# CRISPR-nRAGE, a Cas9 nickase-reverse transcriptase assisted versatile genetic engineering toolkit for *E. coli*

**DOI:** 10.1101/2020.09.02.279141

**Authors:** Yaojun Tong, Tue S. Jørgensen, Christopher M. Whitford, Tilmann Weber, Sang Yup Lee

**Affiliations:** The Novo Nordisk Foundation Center for Biosustainability, Technical University of Denmark, 2800 Kgs. Lyngby, Denmark; Department of Chemical and Biomolecular Engineering, BioProcess Engineering Research Center, BioInformatics Research Center, Institute for the BioCentury, Korea Advanced Institute of Science and Technology (KAIST), Daejeon 34141, Republic of Korea

**Keywords:** CRISPR prime editing (PE), CRISPR-nRAGE, M-MLV reverse transcriptase, H840A Cas9 nickase (Cas9n), genetic engineering, DSB-free, *E. coli*

## Abstract

In most prokaryotes, missing and poorly active non-homologous end joining (NHEJ) DNA repair pathways heavily restrict the direct application of CRISPR-Cas for DNA double-strand break (DSB)-based genome engineering without providing editing templates. CRISPR base editors, on the other hand, can be directly used for genome engineering in a number of bacteria, including *E. coli*, showing advantages over CRISPR-Cas9, since they do not require DSBs. However, as the current CRISPR base editors can only engineer DNA by A to G or C to T/G/A substitutions, they are incapable of mediating deletions, insertions, and combinations of deletions, insertions and substitutions. To address these challenges, we developed a Cas9 nickase (Cas9n)-reverse transcriptase (Moloney Murine Leukemia Virus, M-MLV) mediated, DSB-free, versatile, and single-nucleotide resolution genetic manipulation toolkit for prokaryotes, termed CRISPR-nRAGE (CRISPR-Cas9n Reverse transcriptase Assisted Genome Engineering) system. CRISPR-nRAGE can be used to introduce substitutions, deletions, insertions, and the combination thereof, both in plasmids and the chromosome of *E. coli*. Notably, small sized-deletion shows better editing efficiency compared to other kinds of DNA engineering. CRISPR-nRAGE has been used to delete and insert DNA fragments up to 97 bp and 33 bp, respectively. Efficiencies, however, drop sharply with the increase of the fragment size. It is not only a useful addition to the genome engineering arsenal for *E. coli*, but also may be the basis for the development of similar toolkits for other organisms.

## Main

Advances in synthetic biology, metabolic engineering, multi-omics, high throughput DNA sequencing and synthesis, and computational biology have prompted a rapidly increasing demand for fast and robust genetic engineering methods to speed up the strain development in a Design-Build-Test-Learn cycle. The classic genetic engineering approaches in prokaryotes often are phage-derived RecET and lambda red recombinase-based recombineering^1, 2^. They employ the homology-directed integration/replacement of a donor double stranded DNA (dsDNA) or oligonucleotide for making insertions, deletions, and substitutions of the target DNA. For example, the “Multiplex Automated Genome Engineering” (MAGE)^3^ is a method that can be used for simultaneous manipulation of genes across multiple chromosomal loci of *E. coli*. Possible mutations include mismatch mutation, insertion, and deletion, and editing efficiencies are usually below 20% for all types of edits. The application of MAGE not only needs the synthesis and delivery of ssDNA oligos but also requires the inactivation of mismatch repair (MMR) pathway and the expression of lambda (λ) red recombinase systems (Exo, Beta and Gam) in the target *E. coli* strain^3^. The requirement for an automated setup in order to achieve optimal results causes an elevated background mutation rate in MMR deficient strains, which in turn causes an accumulation of unwanted mutations with each cycle, as well as decreasing allelic replacement efficiencies for increasing sizes of mutations^3^.

CRISPR-Cas (Clustered regularly interspaced short palindromic repeats-CRISPR associated (Cas) proteins) systems, originating from the bacterial adaptive immune system^4^, have been engineered as genome editing tools for a variety of organisms^5^. Among these tools, the Class 2, type II CRISPR system CRISPR-Cas9 of *Streptococcus pyogenes* has been most widely studied and applied. The Cas9 nuclease can be guided by an engineered RNA (single guide RNA, sgRNA) to make DNA double strand breaks (DSBs) of the protospacer adjacent motif (PAM)-containing target DNA^6^. Different types of genetic engineering can be achieved during the repair of DSBs. There are two major pathways for DSB repair *in vivo*, the non-homologous end joining (NHEJ) and the homology-directed repair (HDR)^7^. In most eukaryotes, NHEJ is the dominant way to repair DSBs. During NHEJ repair, small insertions and/or deletions (indels) are introduced at the lesion site, leading to gene disruptions in the target gene. In most bacteria, DSBs normally lead to cell death due to the lack of NHEJ^8^. In these organisms, DNA damage is primarily repaired via HDR, where a template DNA replaces the damaged DNA fragment by recombination^9^.

The lack of NHEJ repair in most prokaryotes restricts the direct use of CRISPR-Cas9 without providing editing templates as a genome editing tool. However, the method is widely used for negative selection to eliminate wild-type cells in recombination-based engineering methods^10^. Unlike CRISPR-Cas9, DSB-free CRISPR base editing systems have successfully been applied for direct genome editing in a number of bacteria without providing editing templates^11-13^. As they rely on DNA deaminase reactions, CRISPR base editors can only make one type of changes to the DNA: the substitution (C to T/A/G, or A to G), and the target C or A has to be within the relatively narrow editing window. Hence, it soon becomes a bottleneck of applying CRISPR base editors for bacterial genome engineering.

Recently, reverse transcriptase-Cas9 H840A nickase (Cas9n)-mediated targeted prime editing (PE) has been demonstrated in human cells^14^, rice and wheat cells^15^ to directly knock-out, knock-in, and replace nucleotides at the target locus without introducing DSBs as well as requiring editing templates. The CRISPR-PE system uses the 3 prime extension sequence of the modified sgRNA (herein named as PEgRNA) to provide a primer binding sequence (PBS) and a reverse transcription template (RTT) carrying the desired edits for reverse transcription with the reverse transcriptase that is allocated in the target locus by Cas9n:sgRNA. After DNA repair, designed mutations are introduced into the target locus (Fig 1d). As the system only introduces a nick in one DNA strand, we hypothesized that it may not cause cell death in bacteria and thus could be applicable in bacterial genome engineering as well. Here, we report the establishment and evaluation of the *E. coli* CRISPR prime editor CRISPR-nRAGE.

**Figure 1.**
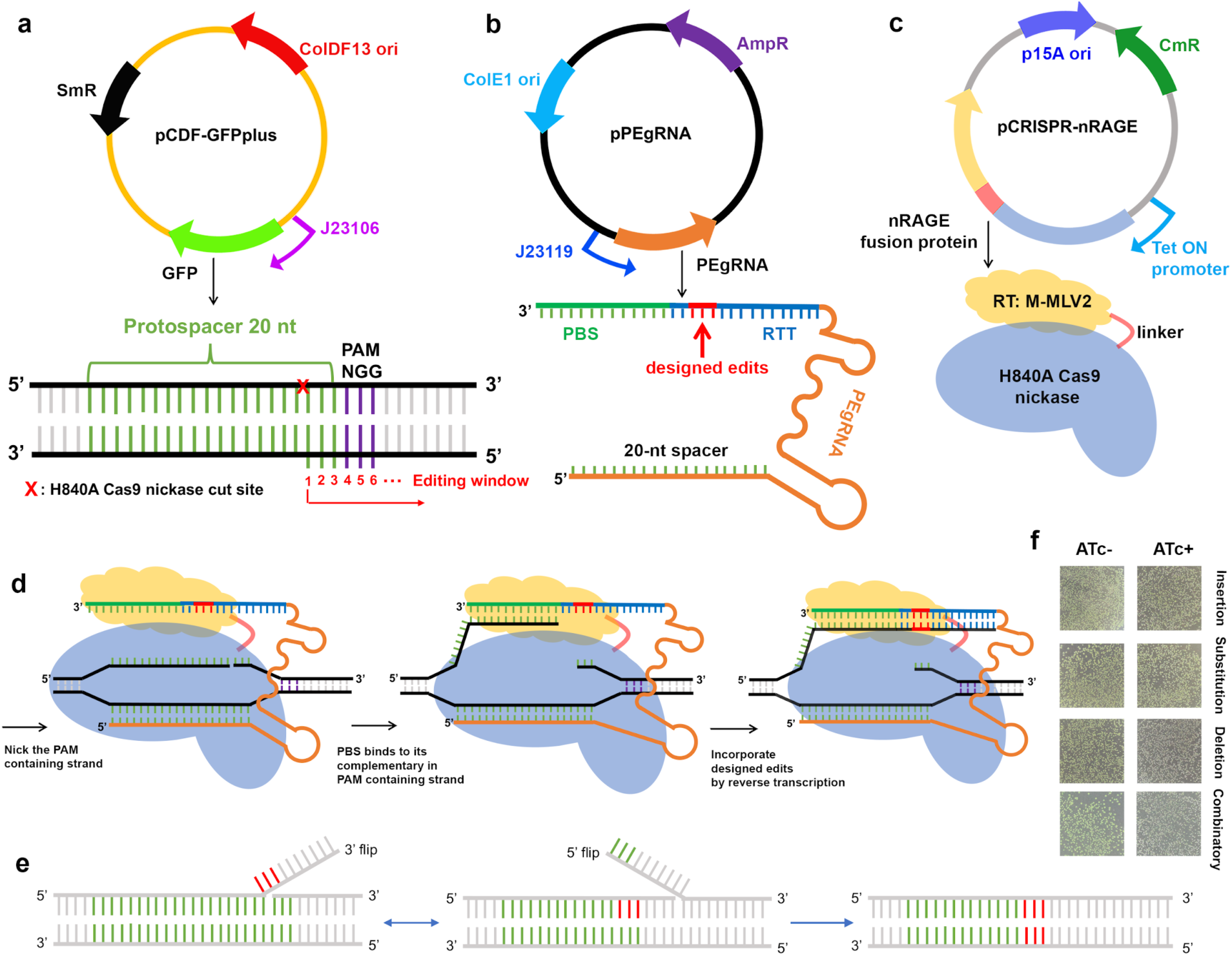
A three-plasmid system for evaluation of CRISPR-nRAGE in *E. coli*. **a**. The plasmid map of the reporter vector pCDF-GFPplus, which carries a constitutive promoter J23106 driving expression of fast folding GFP. The plasmid contains a spectinomycin-resistance (SmR) gene, and the ColDF13 origin (ori). An illustration of the target DNA composition is shown below the plasmid map. **b**. The plasmid map of the PEgRNA transcript bearing vector, which contains the ColE1 ori and an ampicillin-resistance (AmpR) gene for selection. The PEgRNA transcript is under control of the constitutive promoter J23119. A detailed structure is shown beneath, the 3 prime extension sequence is composed of a PBS (in green) and a RTT (in blue), which carries the intended edits (in red). **c**. The plasmid map of the CRISPR-nRAGE vector, which carries a p15A ori and a chloramphenicol-resistance gene (CmR) for selection. The *E. coli* codon optimized, tetracycline inducible promoter driven nRAGE fusion protein consists of a H840A Cas9 nickase (Cas9n), a 33-aa flexible linker, and a moloney murine leukaemia virus (M-MLV) variant M-MLV2, described previously^14^ with the following mutations: D200N, L603W, T306K, W313F, and T330P compared to the WT M-MLV (GenBank: <u>AAC82568.2</u>). **d**. A schematic model for DNA engineering with CRISPR-nRAGE. After being expressed, the Cas9n-M-MLV2:PEgRNA complex binds to the targeted DNA sequence in a sgRNA- and PAM-dependent manner. The Cas9n domain within the fusion protein nicks the PAM-containing strand, freeing the adjacent DNA sequence. Subsequently, this piece of single stranded DNA hybridizes to the PBS, then primes reverse transcription of new DNA containing the designed edits based on the RTT within the 3 prime extension of the PEgRNA transcript. **e**. Two possible consequences after editing by CRISPR-nRAGE. It normally has an equilibration between the edited 3 prime flap and the unedited 5 prime flap, only the cleavage of the 5 prime flap leads to the desired editing. **f**. Colony views of *E. coli* strains transformed with CRISPR-nRAGE systems carrying designed edits of TAA insertion, T to A substitution, T deletion and the combinatorial edits with and without 200 ng/mL ATc induction using a Doc-It imaging station, non-green colonies appeared after 24 h induction of 200 ng/mL ATc.

## Results

### Design of CRISPR-nRAGE for *E. coli*

To evaluate if the reverse transcriptase-Cas9n-mediated DNA modification works in bacteria, we designed a system named CRISPR-Cas9n Reverse transcriptase Assisted Genome Engineering (CRISPR-nRAGE). For testing in *E. coli*, we constructed a three-plasmid system (pCDF-GFPplus, pPEgRNA, and pCRISPR-nRAGE). Plasmid pCDF-GFPplus serves as the reporter plasmid harbouring a gene encoding an *E. coli* codon optimized fast folding GFP^16^ under a constitutive promoter J23106 (Fig. 1a). Plasmid pPEgRNA carries the constitutive promoter J23119 driving PEgRNA transcription. The PEgRNA is composed of a 20-nt spacer and a 3 prime extension containing the PBS and RTT (Fig. 1b). The third plasmid pCRISPR-nRAGE expresses an *E. coli* codon optimized fusion protein composed of an engineered reverse transcriptase M-MLV2 (moloney murine leukaemia virus variant^14^), a flexible linker, and a Cas9n (Cas9 nickase, the H840A mutant of Cas9) under a tetracycline-inducible promoter (Fig. 1c).

### Validation of CRISPR-nRAGE on plasmid editing in *E. coli*

To assess the versatility of the CRISPR-nRAGE system on plasmid DNA engineering in *E. coli*, we designed a full set of possible DNA engineering events, including insertions, deletions, substitutions, and combinations of these to introduce premature stop codons into the coding sequence of GFP. The loss of fluorescence enables easy screening and evaluation for desired editing events. We identified a protospacer located at positions 178-197 of the GFP coding sequence (Supplementary Table 3) that should allow the introduction of a stop codon by DNA engineering with designed PEgRNA. The testing was initiated following observations reported in human cells^14^ with a 3 prime extension consisting of 13 nt PBS and 13 nt RTT scaffold. In the case of insertion, the length of RTT equals the RTT scaffold size plus the designed insertion, for example the length of RTT for TAA insertion is 16 nt (Supplementary Table 3 and Supplementary Table 4). Designed edits were placed inside the potential editing window starting from the nick and continuing downstream^14^ (Fig. 1a). The nRAGE fusion protein binds to the desired PEgRNA transcript, forming an RNA-protein complex, the Cas9n-component of the complex subsequently finds its target DNA sequence and introduces a nick in the PAM containing DNA strand. The PBS within the 3 prime extension then binds to the flipped PAM containing DNA sequence, initiating the reverse transcription to elongate the nicked DNA sequence based on the sequence of the RTT (Fig. 1d). After the reverse transcription process, the nicked double stranded DNA undergoes an equilibration between the edited 3′ flap and the unedited 5′ flap. The cleavage of the unedited 5’ flap then leads to the desired DNA editing^14^ (Fig. 1e).

As a proof of concept, we transformed *E. coli* cells with CRISPR-nRAGE systems programmed for TAA insertion, T to A substitution, T deletion, and the combination thereof. All of these edits will lead to stop codon introduction (Supplementary Table 4). Comparison of GFP expression in 200 ng/mL anhydrotetracycline (ATc) induced and uninduced LB plates, we observed non-fluorescent colonies in all four induced plates (Fig. 1f). In order to further confirm that GFP fluorescence loss was due to the designed DNA editing consequences, we randomly Sanger sequenced eight non-fluorescent colonies from each induced plate. Results demonstrated that almost all of the non-fluorescent colonies were indeed carrying the designed stop codon edits (Fig. 2). Besides the combinatorial editing of insertion, deletion and substitution, we also investigated the possibility of performing double substitutions from one construct. An edit replacing tyrosine at the position 66 of the GFP to histidine (Y66H) was designed by flipping the TAT codon to a CAC codon. We obtained eight non-fluorescent clones out of which 6 clones were correctly edited (Supplementary Fig. 1). However, we also noticed unintended frameshifts in the GFP coding gene in some colonies, resulting in the loss of fluorescence (Supplementary Fig. 2). We classified this as target specific off-target effects, which only happens in a small range around the nick site.

**Figure 2.**
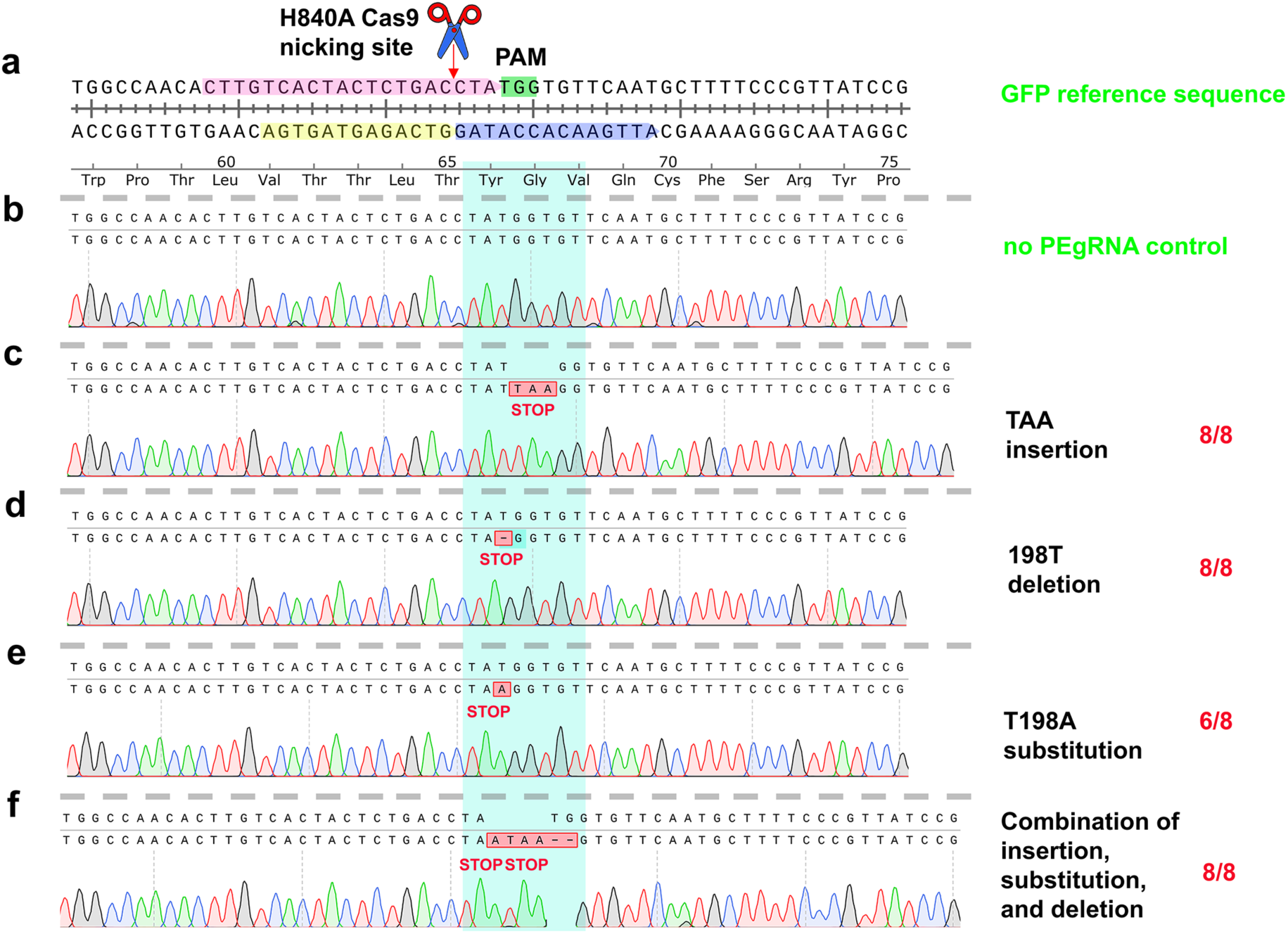
Sanger sequencing traces of four different genetic engineering events achieved by CRISPR-nRAGE. Eight randomly picked colonies of each designed DNA engineering were Sanger sequenced and traces were aligned to the targeted locus of the GFP coding sequence. The correctly edited colony numbers and the total sequenced numbers are shown in red on the right of the figure. **a**. Shown is the target locus of GFP and the potential Cas9 H840A nicking site is indicated by a red arrow. The 20 nt protospacer, the PAM sequence, the PBS, and the RTT are highlighted in pink, green, yellow, and blue, respectively. The translated amino acid sequences are shown underneath each nucleotide sequence. **b**. A non-edited clone served as a negative control. **c**. to **f**. Sanger sequencing traces from edited clones of TAA insertion, a T deletion, T to A substitution, and the combination of insertion, deletion, and substitution, respectively. Introduced stop codons are labeled under the nucleotide sequence, and the potential editing position was highlighted by a cyan box. Numbers in red on the right side show the correct and total Sanger sequenced colonies.

### Characterization of CRISPR-nRAGE in *E. coli*

As the nRAGE fusion protein is driven by the ATc inducible promoter, we evaluated the optimal condition of induction using eight different ATc concentrations. The editing efficiencies were defined by calculating the ratio of non-green colonies. We observed a dose dependent induction manner for all four designed DNA engineering events (Fig. 3a). For cases of 1-bp deletion, 3-bp insertion, 1-bp substitution, and the combinatorial editing, CRISPR-nRAGE can reach efficiencies up to 43.7%, 13.8%, 19.9%, and 2.1%, respectively with 1000 ng/mL of inducer (Fig. 3a). Deletion and insertion of similar sized DNA fragments has an efficiency equal to, or higher than, MAGE^3^. Among the four DNA editing events, deletion showed the highest, while substitution showed the lowest efficiency (Fig. 3a). The editing efficiency did not change significantly with ATc concentrations of 100 ng/mL to 500 ng/mL, and thus we decided to induce CRISPR-nRAGE with 200 ng/mL ATc in the following experiments unless specified otherwise.

**Figure 3.**
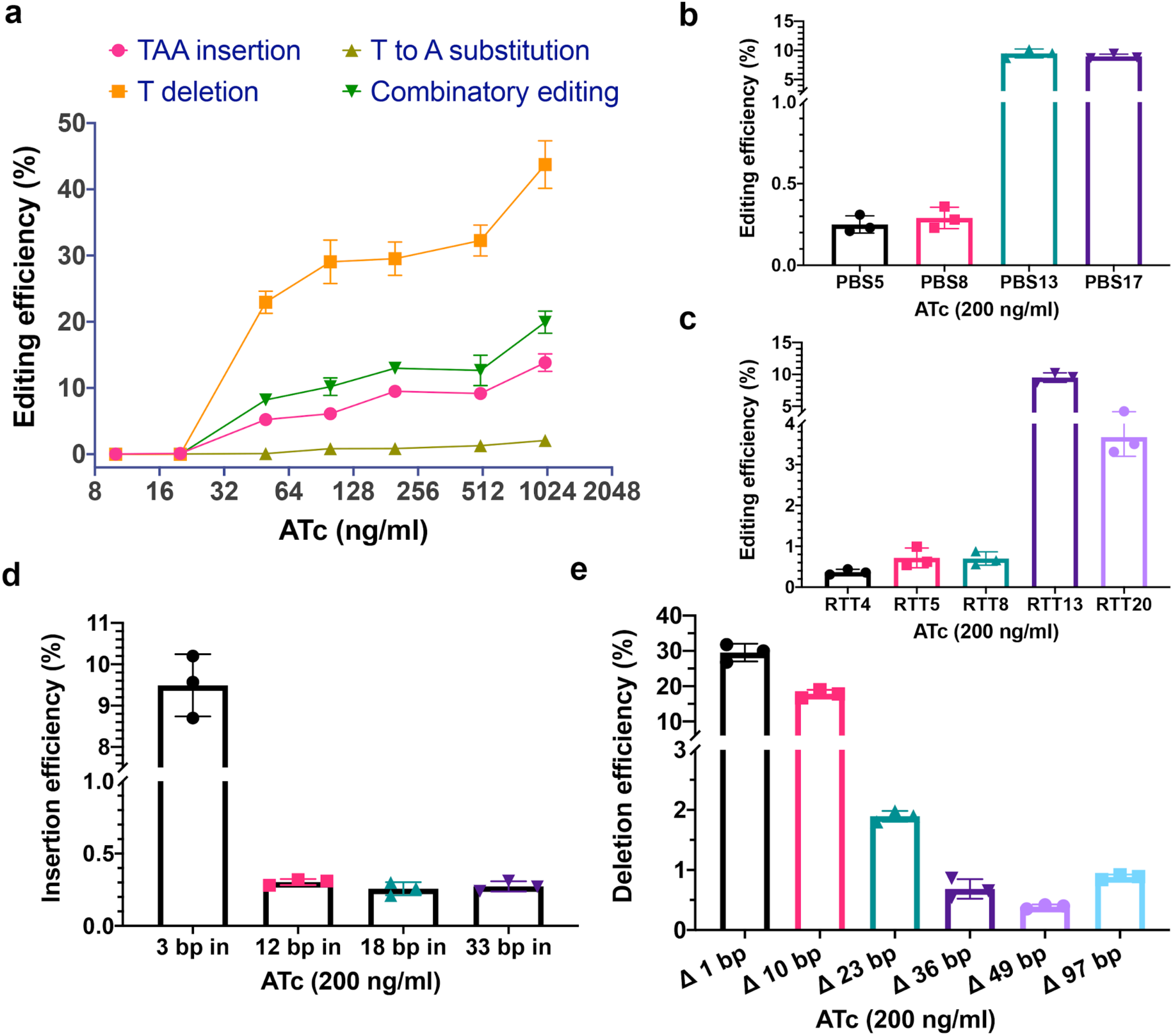
Characteristics of CRISPR-nRAGE for DNA engineering in *E. coli*. **a**. Eight different concentrations of ATc, ranging from 0 ng/mL to 1000 ng/mL (0, 10, 20, 50, 100, 200, 500, and 1000) were used to evaluate the induction of CRISPR-nRAGE on four DNA engineering events of 3-bp insertion, 1-bp deletion, 1-bp substitution, and the combinatorial editing. **b**. The evaluation of PBS length. **c**. The evaluation of RTT scaffold length. **d**. The capacity of DNA fragment insertion with different sizes. **e**. The capacity of DNA fragment deletion with different sizes. Mean ± s.d. of three biological replicates are shown. 200 ng/ml of ATc was used for **b**. to **e**.

Next, we evaluated the optimal length of PBS and RTT by measuring the frequency of the TAA-STOP codon insertion. As a 20-nt spacer was used in the sgRNA construct, the maximum theoretical length of PBS is 17 nt. Nearly no edits were observed when a PBS of 5 nt or 8 nt was used, while a 17 nt PBS showed editing efficiency equivalent to a 13 nt PBS (Fig. 3b). This indicated that the window of PBS for CRISPR-nRAGE is 13 nt to 17 nt. For the RTT scaffold, we designed five different lengths, 4, 5, 8, 13, and 20 nt. We observed that too short or too long RTT reduced the editing efficiency, and the optimal length of the RTT scaffold was around 13 nt (Fig. 3c, and 3d). Results obtained in this study are consistent with previous reports in eukaryotes^14, 15^ Moving forward, by using the optimal length of PBS and RTT scaffold, we systematically tested the capacity of both insertion and deletion with 200 ng/mL of inducer. We designed insertions of 3, 12, 18, and 33 bp, in which 18 bp fragment is the mini-T7 promoter; and deletions of 1, 10, 23, 36, 49, and 97 bp. Though clones with all designed DNA engineering events could be successfully obtained, the editing efficiency dropped greatly with increasing size (Fig. 3d, and 3e). For instance, under 200 ng/mL ATc, editing efficiencies of 1-bp deletion and 10-bp deletion can reach 29.5% and 17.8%, respectively, while efficiencies dropped to below 2% with lengths of 23 bp-97 bp. For insertion, the efficiency was in general lower than deletion. The efficiency of 3-bp insertion was about 10% with 200 ng/mL ATc, and it dropped to below 1% when the size increased to 13 bp-33 bp (Fig. 3d, and 3e).

To evaluate if a second nick in the complementary strand can increase the editing efficiency of CRISPR-nRAGE in prokaryotes like it is described in some mammalian cells^14^ and plant cells^15^, we designed and validated two strategies of the second nick introduction (Supplementary Note 1). There were almost no visible colonies after the second nick was introduced (Supplementary Fig. 3). This result indicates that for organisms not equipped with the NHEJ pathway, introducing the second nick cannot increase the editing efficiency but only compromises the use of CRISPR-nRAGE.

### Assessing the viability of *E. coli* chromosomal DNA editing with CRISPR-nRAGE

In our proof-of-concept experiments, a plasmid-encoded GFP reporter gene was targeted. Therefore, we also assessed if CRISPR-nRAGE is capable of engineering chromosomal DNA. To this end, two metabolic pathways for lactose and D-galactose degradation in *E. coli* MG1655 were selected. β-galactosidase, encoded by the *lacZ* gene within the lactose metabolic pathway, metabolizes X-gal (5-bromo-4-chloro-3-indolyl-β-D-galactoside, an analog of lactose) into 5-bromo-4-chloro-indoxyl, which will form dark blue 5,5’-dibromo-4,4’-dichloro-indigo by oxidation. On the contrary, X-gal remains colorless if the *lacZ* gene is inactivated (Fig. 4a and 4b). An early stop codon was designed to be introduced into the *lacZ* gene of *E. coli* MG1655 by insertion of TAG, deletion of GC, and substitution of GT to TA. In general, the editing efficiencies similar to those for plasmid DNA engineering were observed by counting the white colonies out of the total formed colonies on X-gal supplemented LB plates (Fig. 4c). Editing efficiencies of substitution, insertion and deletion were 6.8%, 12.2%, and 26%, respectively. To validate the editing events, the targeted region of eight randomly picked non-blue colonies from each designed DNA engineering event were PCR amplified and further subjected to Sanger sequencing. All sequenced clones bore the expected edits (Fig. 4d-4f). Moreover, another gene, the *galK* gene from the Leloir pathway of D-galactose metabolism in *E. coli* MG1655 was tested. The loss of function of the *galK* gene can be positively selected by supplementing a galactose analogue 2-deoxy-D-galactose (2-DOG), as 2-DOG will be metabolized by galactokinase (encoded by the *galK* gene) to form a toxic compound 2-deoxy-galactose-1-phosphate, which cannot be further metabolized^17^ (Fig. 4g and 4h). A TAA stop codon was designed to be inserted into the *galK* gene in *E. coli* MG1655 strain. Of the visible colonies on the 2-DOG supplemented M63 agar plate, all four that were randomly picked showed the expected insertion (Fig. 4i).

**Figure 4.**
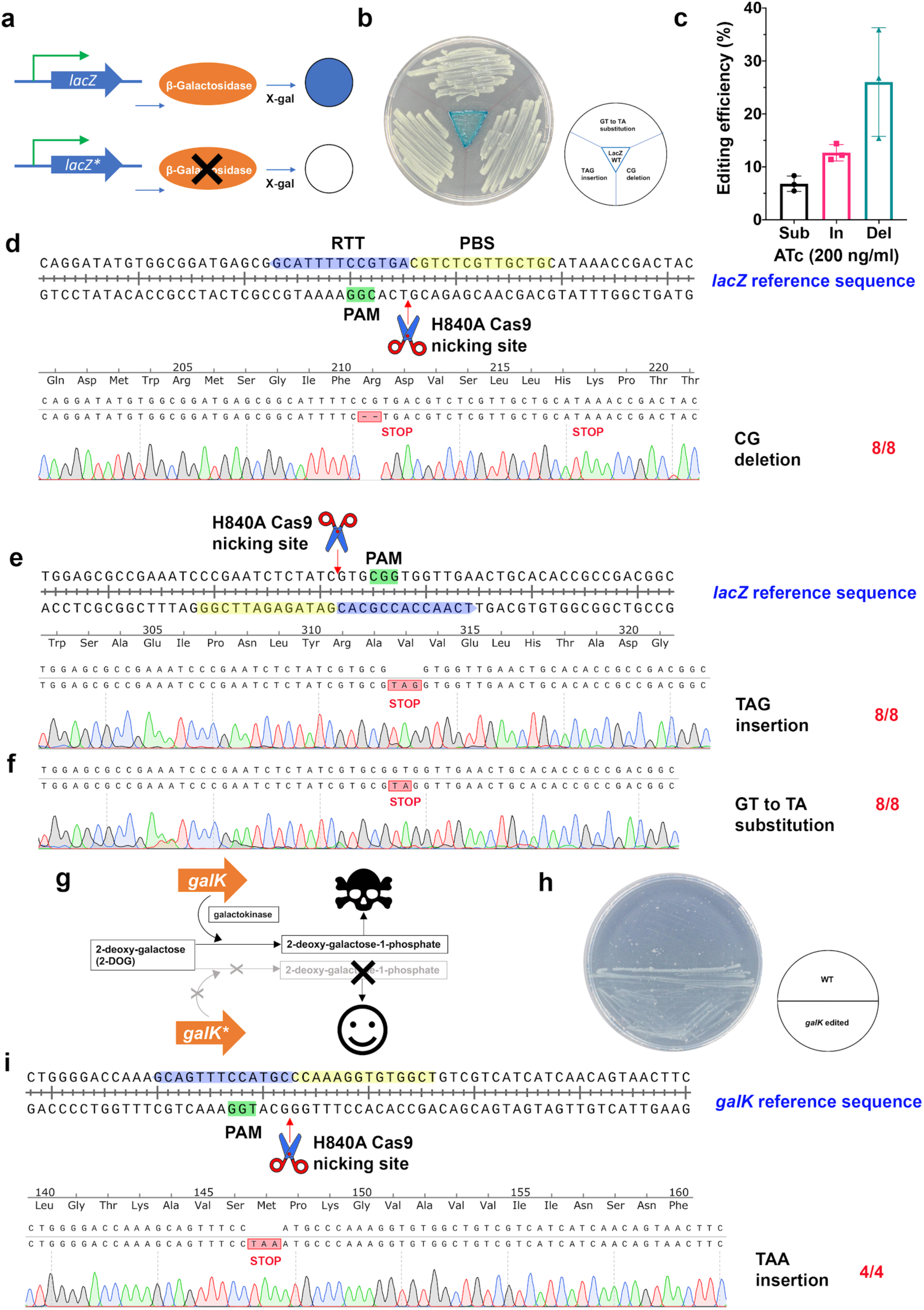
CRISPR-nRAGE is capable of chromosomal DNA engineering. **a**. A graphic illustration of the function of *lacZ* gene. The star within the gene box represents a stop codon being introduced. **b**. An agar plate view of the successful *lacZ* gene inactivation in *E. coli* MG1655 strains by CRISPR-nRAGE. **c**. A bar chart shows the editing efficiencies of chromosomal DNA engineering of 3-bp insertion, 2-bp deletion and 2-bp substitution by CRISPR-nRAGE. **d**. to **f**. Eight non-blue colonies were picked and Sanger sequenced. Sequencing traces aligned with non-edited *lacZ* reference sequences are displayed. Shown are alignments of 2-bp deletion (**d**.); 3-bp (stop codon TAG) insertion (**e**.); and 2-bp substitution (**f**.). **g**. A graphic illustration of the function of *galK* gene. **h**. An agar plate view of the successful g*alK* gene inactivation in *E. coli* MG1655 strains by CRISPR-nRAGE. **i**. Four non-blue colonies were picked and Sanger sequenced. Sequencing traces aligned with non-edited *galK* reference sequences are displayed. The alignment of 3-bp (stop codon TAA) insertion is shown. The potential Cas9 H840A nicking site is indicated by a red arrow. The 20 nt protospacer, the PAM sequence, the PBS, and the RTT are highlighted in pink, green, yellow, and blue, respectively. The translated amino acid sequences together with the introduced stop codons are labeled underneath each nucleotide sequence. Numbers in red on the right side show the correct and total Sanger sequenced colonies.

## Discussion

Due to the importance of *E. coli* in basic microbiological studies and biotechnological applications as a workhorse for the production of various bioproducts, there has been continued demand for novel and efficient DNA engineering tools. Like most prokaryotes, *E. coli* has no functional NHEJ pathway to repair fatal DNA damages like DSBs. Instead, *E. coli* primarily employs homologous recombination-based repair, which made the RecET and lambda red recombineering the major method of DNA engineering^1, 2^. Although much effort has been exerted to simplify and improve the recombineering protocol, it is either still relatively difficult to operate^1, 18^, and it requires the target strain to have a specific genetic background, for example the deficiency of methyl-directed mismatch repair or RecA, the key enzyme for recombinational DNA repair^19^. The emergence of CRISPR-based genetic engineering has been revolutionizing biotechnology, however much less applications were reported in prokaryotes than in eukaryotes^5^, partially because of the different dominant DSB repair pathways. As a result, in many bacteria, including *E. coli*, CRISPR-Cas9 has been widely employed as a tool for counter-selection to eliminate non-modified cells from a mixed population for enrichment of gene-modified cells by homology-directed recombination methods such as the lambda red recombination systems^20, 21^. It remains very challenging to engineer DNA at a single nucleotide level, even when combined with a powerful counter-selection system such as CRISPR-Cas9, the efficiency of making point mutations using oligonucleotide-directed mutagenesis is very low (< 3% before optimization)^22^.

Recently, two types of CRISPR base editors were developed, which are capable of C to T conversion (CBE) by the cytosine deaminase (APOBEC1 or Target-AID)^23, 24^, C to G or A substitution with engineered cytosine deaminases^25^, and A to G conversion (ABE) by the adenosine deaminase (TadA)^12^ without involving DSBs. Thus, they can be directly applied in bacteria for DNA manipulation. One of the main applications is gene inactivation using CBE to convert Arg, Gln, or Trp codons to a stop codon^11, 13^. There are also a few cases of using adenosine deaminase based base editing for *in vivo* protein engineering^11, 12^. However, so far, CRISPR base editing technology has not been as widely used in bacteria as expected due to the restriction of fixed substitutions (C to T/G/A, or A to G) and the relatively narrow editing window (5-7 nucleotides).

We demonstrated in this study that CRISPR-nRAGE cannot only make substitutions but also insertion, deletion, and combinatorial editing at single base pair resolution in *E. coli* without requiring DSBs, editing templates, or homologous recombination. CRISPR-nRAGE has great potential in expanding the possibilities of DNA engineering, although further studies are required to further increase its editing efficiency. As a result of its high modularity and simple composition, CRISPR-nRAGE might be multiplexed by providing a PEgRNA self-processing machinery like Csy4^11^ and consequently applied for high-throughput mutagenesis applications. CRISPR-nRAGE also has the potential of being applicable to a wider range of bacteria including those previously considered difficult to be genetically engineered. As in the case of CRISPR-Cas9 and CRISPR base editing systems, the use of other Cas proteins and protein engineering will likely improve editing capabilities by expanding the selection of accepted PAMs and increasing efficiencies^26-29^. Different reverse transcriptases other than the M-MLV could also provide different features to increase the performance of CRISPR prime editing systems in applied organisms. Modulating intracellular DNA repair systems and better designed PEgRNAs could also be helpful in increasing the editing efficiency.

CRISPR-nRAGE, a versatile DNA engineering system reported in this study, represents a powerful addition to the toolbox of genetic and metabolic engineers not only for E. coli, but other organisms. These tools are likely to substantially advance our understanding of basic life science and to increase capabilities for advanced microbial engineering for biotechnological purposes.

## Materials and methods

### Stains, plasmids, media and growth condition

All *Escherichia coli* strain and plasmids used in this study are listed in Supplementary Table 1. *E. coli* cultures were grown at 37 °C in LB (both broth and solid) (Sigma, US). Appropriate antibiotics were supplemented with the following working concentrations: spectinomycin (50 µg/mL), carbenicillin or ampicillin (100 µg/mL), chloramphenicol (25 µg/mL) kanamycin (50 µg/mL) and anhydrotetracycline (0 to 1 µg/mL). M63 minimal medium was used for positive selection of *galK* mutants. It is composed of 2 g/L (NH_4_)_2_SO_4_, 13.6 g/L KH_2_PO_4_, 0.5 mg/L FeSO_4_-7H_2_O, 1 mM MgSO_4_, 0.1 mM CaCl_2_ and 10 μg/mL thiamine, 0.2% glycerol and 0.1% 2-deoxy-D-galactose. 2% agar was supplemented when making agar plates. X-gal (5-bromo-4-chloro-3-indolyl-beta-D-galactopyranoside) was used for screening *lacZ* mutants. Prior to use, each LB plate with appropriate antibiotics is plated with 40 µL of 20 mg/mL X-gal. All chemicals involved in this study were from Sigma, US.

### General protocol of DNA manipulation

All primers, important sequences, spacers and 3 prime extensions used in this study are listed in Supplementary Tables 2, 3, and 4, respectively. Standard protocols were used for DNA (plasmids and genomic DNA) purification, PCR, and cloning. PCR was performed using Q5^®^ High-Fidelity 2× Master Mix (New England Biolabs, US). The point mutation in dCas9 to create H840A Cas9n was made using Q5^®^ Site-Directed Mutagenesis Kit (New England Biolabs, US). DNA assembly was mainly done by using NEBuilder^®^ HiFi DNA Assembly Master Mix (New England Biolabs, US) unless specified otherwise. DNA digestion was performed with FastDigest restriction enzymes (Thermo Fisher Scientific, US) unless specified otherwise. NucleoSpin^®^ Gel and PCR Clean-up kit (Macherey-Nagel, Germany) was used for DNA clean-up both from PCR products and agarose gel extracts. NucleoSpin^®^ Plasmid EasyPure Kit (Macherey-Nagel, Germany) was used for plasmid preparation. Sanger sequencing was carried out using Mix2Seq kit (Eurofins Scientific, Luxembourg). DNA fragments were synthesized by Genscript while oligonucleotides were synthesized by IDT (Integrated DNA Technologies, US).

All kits and enzymes were used according to the manufacturers’ recommendations. We diligently followed all waste disposal regulations of our institute, university, and local government when disposing of waste materials.

### Multi-plasmid system design and plasmid construction

All plasmids constructed in this study have been deposited to Addgene, individual Addgene plasmid number are listed below. As plasmids in the same testing system should be compatible with each other, and therefore they must have different origins of replication (ori). For this purpose, a combination of p15A ori, ColE1 ori, and ColDF13 ori was used.

Synthetic constitutive promoters J23119 (BBa_J23119) and J23106 (BBa_J23106), and the ribosome binding site (RBS) BBa_B0034 were obtained from the registry for standard biological parts in the iGEM Parts Registry (http://parts.igem.org/Main_Page).

The construction of GFP-based reporter plasmid: The plasmid was designed *in silico* to carry the GFP expression cassette, which is composed of a constitutive promoter J23106, a RBS BBa_B0034, a fast folding GFP variant GFP+^16^, and a terminator T0. The GFP+ coding sequence was codon optimized to *E. coli*. The whole cassette was synthesized by Genscript, and assembled into the pCDF-1b plasmid (ColDF13 ori, Millipore, US) replacing the MCS region by Gibson Assembly, and ended up with the plasmid pCDF-GFPplus (Addgene #).

The construction of CRISPR-nRAGE plasmid: Firstly, we created pCas9n(H840A) from pdCas9-bacteria (p15A ori, Addgene plasmid #44249)^30^ by site-specific mutation of 10A of dCas9 to 10D using Q5^®^ Site-Directed Mutagenesis Kit (New England Biolabs, US). Secondly, we designed the 33a linker-M-MLV2 cassette *in silico*. Linker sequence: SGGSSGGSSGSETPGTSESATPESSGGSSGGSS. M-MLV2, a moloney murine leukaemia virus (M-MLV) variant from a previous study^14^ with the following mutations D200N, L603W, T306K, W313F, and T330P compared to the WT M-MLV (GenBank: <u>AAC82568.2</u>). Thirdly, the cassette was codon optimized to *E. coli*, synthesized by Genscript, and then assembled into pCas9n(H840A) to replace the stop codon of Cas9n by Gibson Assembly, resulting in the plasmid pCRISPR-nRAGE (Addgene #). The fusion protein (cargo) Cas9n-linker-M-MLV2 is under control by a tetracycline inducible promoter. The construction of PEgRNA transcript carrying plasmid: The empty PEgRNA plasmid was modified from the pgRNA-bacteria (ColE1 ori, Addgene plasmid #44251)^30^ by removing the 20 bp spacer, named as pPEgRNA (Addgene #). For construction of functional pPEgRNA there were three steps: firstly, a spacer and 3 prime extension were designed *in silico*; secondly, amplification of the functional PEGgRNA cassette using the pPEgRNA as a template was performed concurrently with amplifying the PEgRNA backbone fragment using the primer set (PEgRNA backbone_F and PEgRNA backbone_R); lastly, the functional PEgRNA cassette was assembled into the PEgRNA backbone. Sanger sequencing was used for validation. Spacers and 3 prime extensions were designed both manually and using PrimeDesign^31^.

For introducing the second nick, we constructed pnsgRNA (pSC101 ori, kanR) by replacing the sfGFP expression cassette in pVRb20_992 (Addgene plasmid #49714)^32^ with the sgRNA transcript cassette from pPEgRNA. We first amplified the plasmid backbone of pVRb20_992 and the sgRNA cassette with primer sets of pVRb_backbone_F and pVRb_backbone_R, and sgRNA_cassette_F and sgRNA_cassette_R from pVRb20_992 and pPEgRNA, respectively. Then these two fragments were Gibson assembled and later validated by Sanger sequencing, resulting in pnsgRNA plasmid (Addgene #). Spacers for introducing the second nick in the nsgRNA paired with the related PEgRNA were designed using PrimeDesign^31^.

### High throughput electroporation of multiple plasmids

*In vivo* assay of strains carrying multiple plasmids were performed from freshly transformed *E. coli* DH10β strains. A HT Nucleofector™ System (Lonza, Switzerland) together with 96-well Nucleocuvette plates (Lonza, Switzerland) were used for high throughput electroporation. Before electroporation, the 96-well Nucleocuvette plate was transferred from −20 °C to ice for 10 min. 20 µL of electrocompetent DH10β or MG1655 *E. coli* cells with 10% glycerol were added into each desired well, 0.5 µL of each plasmid (about 30 ng) was subsequently added. A total amount of plasmid DNA of <100 ng per transformation normally performed well. The program used in this study is X_bacteria_14, with the code GN-100. After electroporation, 180 µL of fresh LB broth were added into each well. The cultures were then transferred into a 96-deep well plate containing 200 µL of fresh LB broth (making the transformation culture in total 400 µL) for recovery for 1h at 37 °C and 300 rpm.

### Editing efficiency evaluation

50 μL electroporation culture (400 μL in the cases of a second nick is introduced) of each strain was plated onto appropriate antibiotics containing LB agar plates supplemented with and without inducer, respectively. All plates were covered by aluminium foil and incubated at 37 °C for 24 h. After cultivation, total colonies were counted by a Doc-It imaging station (Fisher Scientific, US.) with a trisection protocol. Non-green colonies in each zone of all three zones were further counted with and without a Blue-Light Transilluminator (Safe Imager 2.0, Thermo Fisher Scientific, US). The editing efficiency was calculated as: the number of non-green colonies in each zone/total number of visible colonies in the same zone.

### Editing events confirmation by Sanger sequencing

Eight to twelve primarily identified positive clones of each strain were picked, and inoculated into 5 mL LB broth with proper antibiotics. After overnight (∼16 h) cultivation, cultures were subjected to plasmid isolation using the NucleoSpin^®^ Plasmid EasyPure Kit (Macherey-Nagel, Germany) or colony PCR using Q5^®^ High-Fidelity 2× Master Mix (New England Biolabs, US) if a chromosomal region was targeted. The isolated plasmids and the cleaned PCR products were Sanger sequenced using the Mix2Seq kit (Eurofins Scientific, Luxembourg) with proper primers. The obtained sequence traces were analyzed and visualized using SnapGene (GSL Biotech, US).

## Acknowledgments

We thank Simon Shaw for proofreading the manuscript. We thank Alexandra Hoffmeyer for excellent technical support in Illumina sequencing. This work was supported by grants from the Novo Nordisk Foundation (NNF10CC1016517, NNF15OC0016226, NNF16OC0021746). S.Y.L. was also supported by the Technology Development Program to Solve Climate Changes on Systems Metabolic Engineering for Biorefineries (NRF-2012M1A2A2026556 and NRF-2012M1A2A2026557) from the Ministry of Science and ICT through the National Research Foundation (NRF) of Korea.

## Data availability

All data used in this study can be found in the manuscript and/or in the supplementary information.

## Author contribution

Y.T., T.W., and S.Y.L, conceived and designed the project. Y.T. and C.M.W. carried out the laboratory experiments and analyzed the data. T.S.J. performed computational analysis. Y.T., T.W. and S.Y.L. wrote the manuscript with input from all authors.

## References

1. Costantino, N. & Court, D.L. Enhanced levels of lambda Red-mediated recombinants in mismatch repair mutants. Proc. Natl. Acad. Sci. U. S. A. 100, 15748–15753 (2003).

2. Muyrers, J.P., Zhang, Y., Testa, G. & Stewart, A.F. Rapid modification of bacterial artificial chromosomes by ET-recombination. Nucleic Acids Res. 27, 1555–1557 (1999).

3. Wang, H.H. et al. Programming cells by multiplex genome engineering and accelerated evolution. Nature 460, 894–898 (2009).

4. Barrangou, R. et al. CRISPR Provides Acquired Resistance Against Viruses in Prokaryotes. Science 315, 1709–1712 (2007).

5. Sander, J.D. & Joung, J.K. CRISPR-Cas systems for editing, regulating and targeting genomes. Nat. Biotechnol. 32, 347–355 (2014).

6. Jinek, M. et al. A programmable dual-RNA-guided DNA endonuclease in adaptive bacterial immunity. Science 337, 816–821 (2012).

7. Kanaar, R., Hoeijmakers, J.H.J. & van Gent, D.C. Molecular mechanisms of DNA double-strand break repair. Trends Cell Biol. 8, 483–489 (1998).

8. Cui, L. & Bikard, D. Consequences of Cas9 cleavage in the chromosome of Escherichia coli. Nucleic Acids Res. 44, 4243–4251 (2016).

9. Hsu, P.D., Lander, E.S. & Zhang, F. Development and Applications of CRISPR-Cas9 for Genome Engineering. Cell 157, 1262–1278 (2014).

10. Cho, J.S. et al. CRISPR/Cas9-coupled recombineering for metabolic engineering of Corynebacterium glutamicum. Metab. Eng. 42, 157–167 (2017).

11. Tong, Y. et al. Highly efficient DSB-free base editing for streptomycetes with CRISPR-BEST. Proc. Natl. Acad. Sci. U. S. A. 116, 20366–20375 (2019).

12. Gaudelli, N.M. et al. Programmable base editing of A•T to G•C in genomic DNA without DNA cleavage. Nature 551, 464–471 (2017).

13. Banno, S., Nishida, K., Arazoe, T., Mitsunobu, H. & Kondo, A. Deaminase-mediated multiplex genome editing in Escherichia coli. Nat. Microbiol. 3, 423–429 (2018).

14. Anzalone, A.V. et al. Search-and-replace genome editing without double-strand breaks or donor DNA. Nature 576, 149–157 (2019).

15. Lin, Q. et al. Prime genome editing in rice and wheat. Nat Biotechnol 38, 582–585 (2020).

16. Scholz, O., Thiel, A., Hillen, W. & Niederweis, M. Quantitative analysis of gene expression with an improved green fluorescent protein. Eur. J. Biochem. 267, 1565–1570 (2000).

17. Warming, S., Costantino, N., Court, D.L., Jenkins, N.A. & Copeland, N.G. Simple and highly efficient BAC recombineering using galK selection. Nucleic Acids Res. 33, e36 (2005).

18. Lajoie, M.J., Gregg, C.J., Mosberg, J.A., Washington, G.C. & Church, G.M. Manipulating replisome dynamics to enhance lambda Red-mediated multiplex genome engineering. Nucleic Acids Res. 40, e170–e170 (2012).

19. Wang, J. et al. An improved recombineering approach by adding RecA to lambda Red recombination. Mol. Biotechnol. 32, 43–53 (2006).

20. Jiang, W., Bikard, D., Cox, D., Zhang, F. & Marraffini, L.A. RNA-guided editing of bacterial genomes using CRISPR-Cas systems. Nat. Biotechnol. 31, 233–239 (2013).

21. Pyne, M.E., Moo-Young, M., Chung, D.A. & Chou, C.P. Coupling the CRISPR/Cas9 System with Lambda Red Recombineering Enables Simplified Chromosomal Gene Replacement in Escherichia coli. Appl. Environ. Microbiol. 81, 5103–5114 (2015).

22. Lee, H.J., Kim, H.J. & Lee, S.J. CRISPR-Cas9-mediated pinpoint microbial genome editing aided by target-mismatched sgRNAs. Genome Res. 30, 768–775 (2020).

23. Komor, A.C., Kim, Y.B., Packer, M.S., Zuris, J.A. & Liu, D.R. Programmable editing of a target base in genomic DNA without double-stranded DNA cleavage. Nature 533, 420–424 (2016).

24. Nishida, K. et al. Targeted nucleotide editing using hybrid prokaryotic and vertebrate adaptive immune systems. Science 353, aaf8729 (2016).

25. Zhao, D. et al. Glycosylase base editors enable C-to-A and C-to-G base changes. Nat. Biotechnol. doi: 10.1038/s41587-020-0648-3 (2020).

26. Kim, Y.B. et al. Increasing the genome-targeting scope and precision of base editing with engineered Cas9-cytidine deaminase fusions. Nat. Biotechnol. 35, 371–376 (2017).

27. Mitsunobu, H., Teramoto, J., Nishida, K. & Kondo, A. Beyond Native Cas9: Manipulating Genomic Information and Function. Trends Biotechnol. 35, 983–996 (2017).

28. Walton, R.T., Christie, K.A., Whittaker, M.N. & Kleinstiver, B.P. Unconstrained genome targeting with near-PAMless engineered CRISPR-Cas9 variants. Science 368, 290–296 (2020).

29. Richter, M.F. et al. Phage-assisted evolution of an adenine base editor with improved Cas domain compatibility and activity. Nat. Biotechnol. 38, 883–891 (2020).

30. Qi, L.S. et al. Repurposing CRISPR as an RNA-guided platform for sequence-specific control of gene expression. Cell 152, 1173–1183 (2013).

31. Hsu, J.Y. et al. PrimeDesign software for rapid and simplified design of prime editing guide RNAs. BioRxiv doi: 10.1101/2020.05.04.077750 (2020).

32. Rhodius, V.A. et al. Design of orthogonal genetic switches based on a crosstalk map of σs, anti-σs, and promoters. Mol. Syst. Biol. 9, 702 (2013).

