## Supporting Information for "CRISPR-nRAGE, a Cas9 nickase-reverse transcriptase assisted versatile genetic engineering toolkit for *E. coli*"

This file contains:

Supplementary note 1

Supplementary Tables 1-4

Supplementary Figures 1-3

### Supplementary note 1

### A second nick does not increase the editing efficiency of CRISPR-nRAGE in *E. coli*

It has been reported that the editing efficiency of the reverse transcriptase-Cas9 H840A nickase (Cas9n)-mediated targeted prime editing in some mammalian cells[1] and plant cells[2] can be increased by introduction of a second nick to the complementary strand within around 100 bp from the site of the PEgRNA-induced nick. However, it also increases NHEJ-mediated indel formation because Cas9 nickase with paired sgRNAs within ~200 bp can introduce targeted DSBs[3, 4]. As most bacteria do not possess a complete NHEJ system[5-7], it makes the DSB a lethal event if no homology-directed repair (HDR) template is provided. It was thus reasoned that it is impossible to apply the strategy of introducing a close nick in the complementary strand in the NHEJ-deficient bacteria, like *E. coli*. To evaluate our hypothesis, we introduced another plasmid to deliver the designed complementary strand nicking sgRNA (nsgRNA). Two approaches for the secondary nick introduction were tested: nicking the non-edited strand within ~200 bp from the first nick (CRISPR-PE3) and nicking the non-edited strand only after the first nicked strand (the complementary strand) has been edited (CRISPR-PE3b). We designed editing events accordingly both in the plasmid-based system and the chromosome. As expected, for all three designed chromosomal DNA engineering and one plasmid DNA deletion of CRISPR-nRAGE, almost no visible colonies were observed after the second nick was introduced (Supplementary Fig. 3).

### Supplementary Table 1. Plasmids and strains involved in this study

| Plasmid | Background | Reference |
|---|---|---|
| pdCas9- bacteria | *E. coli* codon optimized dCas9 is under control by a tetracycline inducible promoter; CamR; p15A ori | Addgene #44249[8] |
| pgRNA-bacteria | sgRNA transcript carrying plasmid, under control by a constitutive promoter J23119; AmpR; ColE1 ori | Addgene #44251[8] |
| pCDF-b1 | SmR; CloDF13 ori | Millipore, US |
| pCDF-GFPplus | A fast folding GFP variant GFP+ is cloned into pCDF-b1 under control by a constitutive promoter | This study |
| pCRISPR-nRAGE | pdCas9- bacteria is used as the backbone. An *E. coli* codon optimized fusion protein of Cas9n-linker-M-MLV2 is cloned into pdCas9- bacteria by replacing the dCas9. The fusion protein is under control by a tetracycline inducible promoter. | This study |
| pPEgRNA | The 20 bp spacer was removed from pgRNA-bacteria. Therefore, this plasmid carries only a sgRNA scaffold without a spacer. | This study |
| pVRb20_992 | KanR; pSC101 ori; carrying a cassette of Pecf20_992-sfGFP | Addgene #49714[9] |
| pnsgRNA | An sgRNA transcript cassette from pgRNA-bacteria was inserted to replace the Pecf20_992-sfGFP cassette | This study |
| pPEgRNA_GFP_TAA in | pPEgRNA carries a 20 nt spacer targeting GFP coding gene, and a 3 prime extension for inserting a stop codon TAA, the length of both the reverse transcription template and the primer binding sequence is 13 bp | This study |
| pPEgRNA_GFP_TAA in_T4 | The same as pPEgRNA_GFP_TAAin, except that the length of the reverse transcription template is 4 bp | This study |
| pPEgRNA_GFP_TAA in_T5 | The same as pPEgRNA_GFP_TAAin, except that the length of the reverse transcription template is 5 bp | This study |
| pPEgRNA_GFP_TAA in_T8 | The same as pPEgRNA_GFP_TAAin, except that the length of the reverse transcription template is 8 bp | This study |

| | | |
|---|---|---|
| pPEgRNA_GFP_TAAin_T20 | The same as pPEgRNA_GFP_TAAin, except that the length of the reverse transcription template is 20 bp | This study |
| pPEgRNA_GFP_TAAin_PBS5 | The same as pPEgRNA_GFP_TAAin, except that the length of the primer binding sequence is 5 bp | This study |
| pPEgRNA_GFP_TAAin_PBS8 | The same as pPEgRNA_GFP_TAAin, except that the length of the primer binding sequence is 8 bp | This study |
| pPEgRNA_GFP_TAAin_PBS17 | The same as pPEgRNA_GFP_TAAin, except that the length of the primer binding sequence is 17 bp | This study |
| pPEgRNA_GFP_T198A | pPEgRNA carries a 20 nt spacer targeting GFP coding gene, and a 3 prime extension for 198T to 198A substitution within the GFP coding gene, the length of both the reverse transcription template and the primer binding sequence is 13 bp | This study |
| pPEgRNA_GFP_T196C_T198C | pPEgRNA carries a 20 nt spacer targeting GFP coding gene, and a 3 prime extension for 196T , 198T to 196C, 198C substitutions within the GFP coding gene, the length of both the reverse transcription template and the primer binding sequence is 13 bp | This study |
| pPEgRNA_GFP_198Tdel | pPEgRNA carries a 20 nt spacer targeting GFP coding gene, and a 3 prime extension for 198T deletion within the GFP coding gene, the length of both the reverse transcription template and the primer binding sequence is 13 bp | This study |
| pPEgRNA_GFP_combo | pPEgRNA carries a 20 nt spacer targeting GFP coding gene, and a 3 prime extension for 198T to 198A substitution, TAA insertion, and 199G deletion within the GFP coding gene, the length of both the reverse transcription template and the primer binding sequence is 13 bp | This study |
| pPEgRNA_GFP_12in | pPEgRNA carries a 20 nt spacer targeting GFP coding gene, and a 3 prime extension for insertion of a 12 bp fragment within the GFP coding gene, the length of both the reverse transcription template and the primer binding sequence is 13 bp | This study |
| pPEgRNA_GFP_18in | pPEgRNA carries a 20 nt spacer targeting GFP coding gene, and a 3 prime extension for insertion of a 18 bp fragment within the GFP coding gene, the length of both the reverse transcription template and the primer binding sequence is 13 bp | This study |
| pPEgRNA_GFP_33in | pPEgRNA carries a 20 nt spacer targeting GFP coding gene, and a 3 prime extension for insertion of a 33 bp fragment within the GFP coding gene, the length of both | This study |

| | | |
|---|---|---|
| | the reverse transcription template and the primer binding sequence is 13 bp | |
| pPEgRNA_GFP_10del | pPEgRNA carries a 20 nt spacer targeting GFP coding gene, and a 3 prime extension for deletion of a 10 bp fragment within the GFP coding gene, the length of both the reverse transcription template and the primer binding sequence is 13 bp | This study |
| pPEgRNA_GFP_23del | pPEgRNA carries a 20 nt spacer targeting GFP coding gene, and a 3 prime extension for deletion of a 23 bp fragment within the GFP coding gene, the length of both the reverse transcription template and the primer binding sequence is 13 bp | This study |
| pPEgRNA_GFP_36del | pPEgRNA carries a 20 nt spacer targeting GFP coding gene, and a 3 prime extension for deletion of a 36 bp fragment within the GFP coding gene, the length of both the reverse transcription template and the primer binding sequence is 13 bp | This study |
| pPEgRNA_GFP_49del | pPEgRNA carries a 20 nt spacer targeting GFP coding gene, and a 3 prime extension for deletion of a 49 bp fragment within the GFP coding gene, the length of both the reverse transcription template and the primer binding sequence is 13 bp | This study |
| pPEgRNA_GFP_97del | pPEgRNA carries a 20 nt spacer targeting GFP coding gene, and a 3 prime extension for deletion of a 97 bp fragment within the GFP coding gene, the length of both the reverse transcription template and the primer binding sequence is 13 bp | This study |
| pPEgRNA_lacZ_TAGin | pPEgRNA carries a 20 nt spacer targeting *lacZ* gene, and a 3 prime extension for TAG insertion into the *lacZ* gene for a stop codon introduction, the length for the reverse transcription template and the primer binding sequence is 16 bp and 13 bp, respectively | This study |
| pPEgRNA_lacZ_CGdel | pPEgRNA carries a 20 nt spacer targeting *lacZ* gene, and a 3 prime extension for CG deletion in the *lacZ* gene for a stop codon introduction, the length for the reverse transcription template and the primer binding sequence is 18 bp and 14 bp, respectively | This study |
| pPEgRNA_lacZ_GTtoTAsub | pPEgRNA carries a 20 nt spacer targeting *lacZ* gene, and a 3 prime extension for GT to TA substitution in the *lacZ* gene for a stop codon introduction, the length of both the reverse transcription template and the primer binding sequence is 14 bp | This study |
| pPEgRNA_galK_TAAin | pPEgRNA carries a 20 nt spacer targeting *galK* gene, and a 3 prime extension for TAG insertion into the *galK* gene for a stop codon introduction, the length for the reverse transcription template and the primer binding sequence is 16 bp and 13 bp, respectively | This study |

| | | |
|---|---|---|
| pnsgRNA_GFP_TAAin | pnsgRNA carries a 20 nt spacer that pairs with pPEgRNA_GFP_TAAin to introduce the second nick | This study |
| pnsgRNA_lacZ_TAGin | pnsgRNA carries a 20 nt spacer that pairs with pPEgRNA_lacZ_TAGin to introduce the second nick | This study |
| pnsgRNA_lacZ_CGdel | pnsgRNA carries a 20 nt spacer that pairs with pPEgRNA_lacZ_CGdel to introduce the second nick | This study |
| pnsgRNA_lacZ_GTtoTAsub | pnsgRNA carries a 20 nt spacer that pairs with pPEgRNA_lacZ_GTtoTAsub to introduce the second nick | This study |
| **Strain** | **Background** | **Reserence** |
| *Escherichia coli* DH10β (DH10B) | str. K F- mcrA Δ(mrr-hsdRMS-mcrBC) φ80lacZΔM15 ΔlacX74 recA1 endA1 araD139 Δ (ara, leu)7697 galU galK λ- rpsL nupG /pMON14272 / pMON7124 | Thermo Fisher Scientific, US |
| *Escherichia coli* MG1655 | Str. K F-, lambda-, rph-1 | Maintained in lab |
| nRAGE0001 | *E. coli* DH10β carries pCDF-GFPplus | This study |
| nRAGE0002 | *E. coli* DH10β carries pCRISPR-nRAGE | This study |
| nRAGE0003 | *E. coli* DH10β carries pCRISPR-nRAGE and pCDF-GFPplus | This study |
| nRAGE0004 | *E. coli* DH10β carries pCRISPR-nRAGE, pCDF-GFPplus, and pPEgRNA | This study |
| nRAGE0005 | *E. coli* DH10β carries pCRISPR-nRAGE, pCDF-GFPplus, and pPEgRNA_GFP_TAAin | This study |
| nRAGE0006 | *E. coli* DH10β carries pCRISPR-nRAGE, pCDF-GFPplus, and pPEgRNA_GFP_TAAin_T4 | This study |
| nRAGE0007 | *E. coli* DH10β carries pCRISPR-nRAGE, pCDF-GFPplus, and pPEgRNA_GFP_TAAin_T5 | This study |

| | | |
|---|---|---|
| nRAGE0008 | *E. coli* DH10β carries pCRISPR-nRAGE, pCDF-GFPplus, and pPEgRNA_GFP_TAAin_T8 | This study |
| nRAGE0009 | *E. coli* DH10β carries pCRISPR-nRAGE, pCDF-GFPplus, and pPEgRNA_GFP_TAAin_T20 | This study |
| nRAGE0010 | *E. coli* DH10β carries pCRISPR-nRAGE, pCDF-GFPplus, and pPEgRNA_GFP_TAAin_PBS5 | This study |
| nRAGE0011 | *E. coli* DH10β carries pCRISPR-nRAGE, pCDF-GFPplus, and pPEgRNA_GFP_TAAin_PBS8 | This study |
| nRAGE0012 | *E. coli* DH10β carries pCRISPR-nRAGE, pCDF-GFPplus, and pPEgRNA_GFP_TAAin_PBS17 | This study |
| nRAGE0013 | *E. coli* DH10β carries pCRISPR-nRAGE, pCDF-GFPplus, and pPEgRNA_GFP_T198A | This study |
| nRAGE0014 | *E. coli* DH10β carries pCRISPR-nRAGE, pCDF-GFPplus, and pPEgRNA_GFP_T196C_T198C | This study |
| nRAGE0015 | *E. coli* DH10β carries pCRISPR-nRAGE, pCDF-GFPplus, and pPEgRNA_GFP_198Tdel | This study |
| nRAGE0016 | *E. coli* DH10β carries pCRISPR-nRAGE, pCDF-GFPplus, and pPEgRNA_GFP_combo | This study |
| nRAGE0017 | *E. coli* DH10β carries pCRISPR-nRAGE, pCDF-GFPplus, and pPEgRNA_GFP_12in | This study |
| nRAGE0018 | *E. coli* DH10β carries pCRISPR-nRAGE, pCDF-GFPplus, and pPEgRNA_GFP_18in | This study |
| nRAGE0019 | *E. coli* DH10β carries pCRISPR-nRAGE, pCDF-GFPplus, and pPEgRNA_GFP_33in | This study |
| nRAGE0020 | *E. coli* DH10β carries pCRISPR-nRAGE, pCDF-GFPplus, and pPEgRNA_GFP_10del | This study |
| nRAGE0021 | *E. coli* DH10β carries pCRISPR-nRAGE, pCDF-GFPplus, and pPEgRNA_GFP_23del | This study |

| | | |
|---|---|---|
| nRAGE0022 | *E. coli* DH10β carries pCRISPR-nRAGE, pCDF-GFPplus, and pPEgRNA_GFP_36del | This study |
| nRAGE0023 | *E. coli* DH10β carries pCRISPR-nRAGE, pCDF-GFPplus, and pPEgRNA_GFP_49del | This study |
| nRAGE0024 | *E. coli* DH10β carries pCRISPR-nRAGE, pCDF-GFPplus, and pPEgRNA_GFP_97del | This study |
| nRAGE0025 | *E. coli* MG1655 carries pCRISPR-nRAGE and pPEgRNA_lacZ_TAGin | This study |
| nRAGE0026 | *E. coli* MG1655 carries pCRISPR-nRAGE and pPEgRNA_lacZ_CGdel | This study |
| nRAGE0027 | *E. coli* MG1655 carries pCRISPR-nRAGE and pPEgRNA_lacZ_GTtoTAsub | This study |
| nRAGE0028 | *E. coli* MG1655 carries pCRISPR-nRAGE and pPEgRNA_galK_TAAin | This study |

**Supplementary Table 2. Primers used in this study**

| **Names** | **Sequence (5' to 3')** | **Purpose** |
|---|---|---|
| PEgRNA_GFP_F | gtcctaggtataatactagtCTTGTCACTACTCTGACCTAgttttagagctagaaatagc | For PCR amplification of desired PEgRNA functional cassettes. Sequences in lower case are overhangs for Gibson assembly. |
| PEgRNA_GFP_TAAin_R | gggcccaagcttcaaaaaaaTCACTACTCTGACCTATTAAGGTGTTCAAgcaccgactcggtgccactt | |
| PEgRNA_GFP_TAAin_T4_R | gggcccaagcttcaaaaaaaTCACTACTCTGACCTATTAAgcaccgactcggtgccactt | |
| PEgRNA_GFP_TAAin_T5_R | gggcccaagcttcaaaaaaaTCACTACTCTGACCTATTAAGgcaccgactcggtgccactt | |

| | |
|---|---|
| PEgRNA_GFP_TAAin_T8_R | gggcccaagcttcaaaaaaTCACTACTCTGACCTATTAAGGTGgcaccgactcggtgccactt |
| PEgRNA_GFP_TAAin_T20_R | gggcccaagcttcaaaaaaTCACTACTCTGACCTATTAAGGTGTTCAATGCTTTTgcaccgactcggtgccactt |
| PEgRNA_GFP_TAAin_PBS5_R | gggcccaagcttcaaaaaaCTGACCTATTAAGGTGTTCAAgcaccgactcggtgccactt |
| PEgRNA_GFP_TAAin_PBS8_R | gggcccaagcttcaaaaaaACTCTGACCTATTAAGGTGTTCAAgcaccgactcggtgccactt |
| PEgRNA_GFP_TAAin_PBS17_R | gggcccaagcttcaaaaaaACTCTGACCTATTAAGGTGTTCAAgcaccgactcggtgccactt |
| PEgRNA_GFP_T198A_R | gggcccaagcttcaaaaaaTCACTACTCTGACCTAAGGTGTTCAAgcaccgactcggtgccactt |
| PEgRNA_GFP_T196C_T198C_R | gggcccaagcttcaaaaaaTCACTACTCTGACCCACGGTGTTCAAgcaccgactcggtgccactt |
| PEgRNA_GFP_198Tdel_R | gggcccaagcttcaaaaaaTCACTACTCTGACCTAGGTGTTCAATgcaccgactcggtgccactt |
| PEgRNA_GFP_combo_R | gggcccaagcttcaaaaaaTCACTACTCTGACCTAATAAGTGTTCAAgcaccgactcggtgccactt |
| PEgRNA_GFP_12in_R | gggcccaagcttcaaaaaaTCACTACTCTGACCTATGTAATCTGTACAGGTGTTCAAgcaccgactcggtgccactt |
| PEgRNA_GFP_18in_R | gggcccaagcttcaaaaaaTCACTACTCTGACCTATTAATACGACTCACTATAGGGTGTTCAAgcaccgactcggtgccactt |
| PEgRNA_GFP_33in_R | gggcccaagcttcaaaaaaTCACTACTCTGACCTATTGTAGAATCAGCCCACGAAACCGAGCGGGCGAGGGTGTTCAAgcaccgactcggtgccactt |
| PEgRNA_GFP_10del_R | gggcccaagcttcaaaaaaTCACTACTCTGACATATCTTTCAAAGgcaccgactcggtgccactt |

| | | |
|---|---|---|
| PEgRNA_GFP_23del_R | gggcccaagcttcaaaaaaTCACTACTCTGACATATCTTTCAAAG gcaccgactcggtgccactt | |
| PEgRNA_GFP_36del_R | gggcccaagcttcaaaaaaTCACTACTCTGACATATCTTTCAAAG gcaccgactcggtgccactt | |
| PEgRNA_GFP_49del_R | gggcccaagcttcaaaaaaTCACTACTCTGACATATCTTTCAAAG gcaccgactcggtgccactt | |
| PEgRNA_GFP_97del_R | gggcccaagcttcaaaaaaTCACTACTCTGACATATCTTTCAAAG gcaccgactcggtgccactt | |
| pPEgRNA_lacZ_TAGin_F | gtcctaggtataatactagtAATCCCGAATCTCTATCGTGgttttagagctagaaatagc | For PCR amplification of desired functional PEgRNA_lacZ_TAGinca ssettes. Sequences in lower case are overhangs for Gibson assembly. |
| pPEgRNA_lacZ_TAGin_R | gggcccaagcttcaaaaaaCCGAATCTCTATCGTGCGTAGGTGGTTGA gcaccgactcggtgccactt | |
| pPEgRNA_lacZ_CGdel_F | gtcctaggtataatactagtTATGCAGCAACGAGACGTCAgttttagagctagaaatagc | For PCR amplification of desired functional PEgRNA_lacZ_CGde cassettes. Sequences in lower case are overhangs for Gibson assembly. |
| pPEgRNA_lacZ_CGdel_R | gggcccaagcttcaaaaaaGCAGCAACGAGACGTCAGAAAATGCCGCTCATgcaccgactcggtgccactt | |
| pPEgRNA_lacZ_GTtoTAsub_F | gtcctaggtataatactagtGCGAGTTGCGTGACTACCTAgttttagagctagaaatagc | For PCR amplification of desired functional PEgRNA_lacZ_GTtoTA sub cassettes. Sequences in lower case are overhangs |
| pPEgRNA_lacZ_GTtoTAsub_R | gggcccaagcttcaaaaaaAGTTGCGTGACTACTAACGGGTAACAGTgcaccgactcggtgccactt | |

| | | |
|---|---|---|
| | | for Gibson assembly. |
| pPEgRNA_galK_TAAin_F | gtcctaggtataatactagtGACAGCCACACCTTTGGGCAgttttagagctagaaatagc | For PCR amplification of desired functional PEgRNA_galK_TAAin cassettes. Sequences in lower case are overhangs for Gibson assembly. |
| pPEgRNA_galK_TAAin_R | gggcccaagcttcaaaaaaGCCACACCTTTGGGCATTTAGGAAACTGCgcaccgactcggtgccactt | |
| lacZ_check_F | gatgaaagctggctacagga | For PCR amplification of the targeted region in *lacZ* gene. The forward primer is also used for Sanger sequencing. |
| lacZ_check_F | tgacggttaacgcctcgaat | |
| galK_check_F | caatgggctaactacgttcg | For PCR amplification of the targeted region in *galK* gene. The forward primer is also used for Sanger sequencing. |
| galK_check_F | gtcgccaatcacagctttga | |
| pnsgRNA_GFP_TAAin_F | AAAGCATTGAACACCttaATGTTTTAGAGCTAGAAATAGC | For PCR amplification of the nsgRNA_GFP_TAAin cassette. |
| pnsgRNA_GFP_TAAin_R | ATtaaGGTGTTCAATGCTTTactagtattatacctaggac | |
| pnsgRNA_lacZ_TAGin_F | CctaCGCACGATAGAGATTCGTTTTAGAGCTAGAAATAGC | For PCR amplification of the |

| | | |
|---|---|---|
| pnsgRNA_lacZ_TAGin_R | GAATCTCTATCGTGCGtagGactagtattatacctaggac | nsgRNA_lacZ_TAGin cassette. |
| pnsgRNA_lacZ_CGdel_F | CTGGAGTGACGGCAGTTATCGTTTTAGAGCTAGAAATAGC | For PCR amplification of the nsgRNA_lacZ_CGdel cassette. |
| pnsgRNA_lacZ_CGdel_R | GATAACTGCCGTCACTCCAGactagtattatacctaggac | |
| pnsgRNA_lacZ_GTtoTAsub_F | CATTAAAGCGAGTGGCAACAGTTTTAGAGCTAGAAATAGC | For PCR amplification of the nsgRNA_lacZ_GTtoTAsub cassette. |
| pnsgRNA_lacZ_GTtoTAsub_R | TGTTGCCACTCGCTTTAATGactagtattatacctaggac | |
| pVRb_backbone_F | CTGTTGTTTGTCGGTGAACG | For PCR amplification of the pVRb backbone from pVRb_20_992. |
| pVRb_backbone_R | AAGGGCCTCGTGATACGCCT | |
| sgRNA_cassette_F | AGGCGTATCACGAGGCCCTTgaattctaaagatctttgac | For PCR amplification of the sgRNA cassette from pPEgRNA. |
| sgRNA_cassette_R | CGTTCACCGACAAACAACAGataaaacgaaaggcccagtc | |
| nsgRNA_seq | GCAATTCCGACGTCTAAG | For validation of the Gibson assembly of pnsgRNA. Also for validation of 20-bp spacer cloning. |
| PEgRNA_backbone_F | TTTTTTTGAAGCTTGGGCCC | For PCR amplification of the |

| | | |
|---|---|---|
| PEgRNA_backbone_R | ACTAGTATTATACCTAGGAC | pPEgRNA backbone |
| PEgRNA_F | GTCCTAGGTATAATACTAGTGTTTTAGAGCTAGAAATAGCA | For removal of the 20 bp spacer from the plasmid pgRNA-bacteria. |
| PEgRNA_R | ACTAGTATTATACCTAGGACTGAGCTAGCT | |
| PEgRNA_Seq | AATAGGCGTATCACGAGGCA | For Sanger sequencing of correctly assembled PEgRNA plasmids. |
| pCDF-GFP_Seq | GAAATACTAGATGAGCAAAGGAGAAG | For screening of edited events using Sanger sequencing. |
| pCDF-1b_F | GTATATCTCCTTATTAAAGT | For PCR amplification of the pCDF-1b backbone. |
| pCDF-1b_R | ATTAACCTAGGCTGCTGCCA | |
| GFPplus_F | ACTTTAATAAGGAGATATACTTTACGGCTAGCTCAGTCCT | For PCR amplification of the J23106-GFPplus-T0 cassette. |
| GFPplus_R | TGGCAGCAGCCTAGGTTAATCGAACCGAACAGGCTTATGT | |
| GFPplus_check_F | ATGCGACTCCTGCATTAGGA | For screening of correctly assembled pCDF-GFPplus using PCR. The GFPplus_ch |
| GFPplus_check_R | ACTAGTCGCCAGGGTTTTCC | |

| | | |
|---|---|---|
| | | eck_F is also used for Sanger sequencing validation. |
| A10D_F | TAGGCTTAGATATCGGCACAAA | For site-directed mutating of 10A to 10D with the dCas9 in the pdCas9-bacteria. |
| A10D_R | TTGAGTATTTCTTATCCATATG | |
| D10_Seq | GCGAGTTTACGGGTTGTTA | For screening of correct mutations of 10A to 10D using Sanger sequencing. |
| pCas9n_F | TAACTCGAGTAAGGATCTCC | For PCR amplification of the pCas9n(H840A) backbone, the stop codon of Cas9n is removed. |
| pCas9n_R | GTCACCTCCTAGCTGACTCA | |
| EcMMLV2_F | TGAGTCAGCTAGGAGGTGAC | For PCR amplification of the *E. coli* codon optimized linker-M-MLV2 fragment. |
| EcMMLV2_R | GGAGATCCTTACTCGAGTTA | |
| EcMMLV2_check_R | AACAAACAGCTCGAACGGCT | Together with EcMMLV2_F, this primer set is used for PCR screening of *E. coli* codon optimized linker-M-MLV2 |

| | | |
|---|---|---|
| | | fragment insertion. |
| EcMMLV2_seq | AACTGGATTGCCAACAGGGT | Together with EcMMLV2_F, these three primers are used to confirm the insertion of *E. coli* codon optimized linker-M-MLV2 fragment by Sanger sequencing. |
| dblTerm_R_seq | GAAGGTGAGCCAGTGTGACT | |

**Supplementary Table 3. Important sequences involved in this study**

| Name | Sequence | Description |
|---|---|---|
| J23106-GFPplus-T0 cassette | TTTACGGCTAGCTCAGTCCT<br>AGGTATAGTGCTAGCTACTA<br>GAGAAAGAGGAGAAATACTA<br>GATGAGCAAAGGAGAAGAA<br>CTTTTCACTGGAGTTGTCCC<br>AATTCTTGTTGAATTAGATG<br>GTGATGTTAATGGGCACAAA<br>TTTTCTGTCAGTGGAGAGGG<br>TGAAGGTGATGCTACATACG<br>GAAAACTCACCCTTAAATTT<br>ATTTGCACTACTGGAAAACT<br>ACCTGTTCCATGGCCAACAC<br>TTGTCACTACTCTGACCTAT<br>GGTGTTCAATGCTTTTCCCG<br>TTATCCGGATCACATGAAAC<br>GGCATGACTTTTTCAAGAGT<br>GCCATGCCCGAAGGTTATGT<br>ACAGGAACGCACTATATCTT<br>TCAAAGATGACGGGAACTAC<br>AAGACGCGTGCTGAAGTCA<br>AGTTTGAAGGTGATACCCTT<br>GTTAATCGTATCGAGTTAAA<br>GGGTATTGATTTTAAAGAAG<br>ATGGAAACATTCTCGGACAC<br>AAACTAGAGTACAACTATAA | A fast folding GFP variant GFP+ encoding gene (in blue), controlled by a constitutive promoter J23106 (in green) and ended with a T0 terminator (in orange). |

| | | |
|---|---|---|
| | CTCACACAATGTATACATCA<br>CGGCAGACAAACAAAAGAAT<br>GGAATCAAAGCTAACTTCAA<br>AATTCGCCACAACATTGAAG<br>ATGGTTCCGTTCAACTAGCA<br>GACCATTATCAACAAAATAC<br>TCCAATTGGCGATGGCCCT<br>GTCCTTTTACCAGACAACCA<br>TTACCTGTCGACACAATCTG<br>CCCTTTCGAAAGATCCCAAC<br>GAAAAGCGTGACCACATGG<br>TCCTTCTTGAGTTTGTAACT<br>GCTGCTGGGATTACACATG<br>GCATGGATGAGCTCTACAAA<br>TGAAGCGCATACCTGCAGG<br>CATGCAAGCTTGCGGCCGC<br>GTCGTGACTGGGAAAACCC<br>TGGCGACTAGTCTTGGACTC<br>CTGTTGATAGATCCAGTAAT<br>GACCTCAGAACTCCATCTGG<br>ATTTGTTCAGAACGCTCGGT<br>TGCCGCCGGGCGTTTTTTAT<br>TGGTGAGAATCCAGGGGTC<br>CCCAATAATTACGATTTAAAT<br>TTGACATAAGCCTGTTCGGT<br>TCG | |
| *E. coli* codon optimized linker-M-MLV2 fragment | *TGAGTCAGCTAGGAGGTGA*<br>*C*AGCGGCGGCAGCAGCGG<br>CGGCAGCAGCGGCAGCGAA<br>ACCCCGGGCACCAGCGAAA<br>GCGCGACCCCGGAAAGCAG<br>CGGCGGCAGCAGCGGCGG<br>TAGCAGCACCCTGAACATCG<br>AGGACGAGTATCGTCTGCAT<br>GAGACCAGCAAGGAGCCGG<br>ATGTTAGCCTGGGTAGCACC<br>TGGCTGAGCGACTTTCCGC<br>AGGCGTGGGCGGAAACCGG<br>CGGCATGGGTCTGGCGGTT<br>CGCCAGGCGCCGCTGATCA<br>TTCCGCTGAAGGCGACCAG<br>CACCCCGGTTAGCATCAAG<br>CAGTATCCGATGAGCCAGG<br>AAGCGCGTCTGGGTATTAAG<br>CCGCACATTCAACGTCTGCT<br>GGACCAGGGTATTCTGGTG<br>CCGTGCCAGAGCCCGTGGA<br>ATACCCCGCTGCTGCCGGT<br>GAAGAAACCGGGTACCAAT<br>GATTACCGTCCGGTGCAAG<br>ACCTGCGTGAGGTTAACAAG | A 33-amino acid linker (in red) fused with the M-MLV2 encoding sequence (in black). 20 bp overhangs for Gibson assembly into pCas9n(H840A) is shown in gray italics. |

| | |
|---|---|
| | CGCGTTGAAGATATTCATCC GACCGTTCCGAACCCGTAC AACCTGCTGAGCGGTCTGC CGCCGAGCCACCAGTGGTA TACCGTGCTGGATCTGAAG GACGCGTTTTTCTGCCTGCG TCTGCACCCGACCAGCCAA CCGCTGTTCGCGTTTGAATG GCGTGACCCGGAAATGGGT ATCAGCGGCCAACTGACCT GGACCCGTCTGCCGCAGGG CTTTAAAAACAGCCCGACCC TGTTCAACGAGGCGCTGCA CCGTGATCTGGCGGACTTC CGTATCCAACACCCGGATCT GATCCTGCTGCAGTACGTG GACGATCTGCTGCTGGCGG CGACCAGCGAACTGGATTG CCAACAGGGTACCCGTGCG CTGCTGCAGACCCTGGGTA ACCTGGGTTACCGTGCGAG CGCGAAAAAGGCGCAAATTT GCCAGAAGCAAGTGAAGTAT CTGGGCTACCTGCTGAAGG AAGGTCAACGCTGGCTGAC CGAGGCGCGTAAGGAAACC GTTATGGGTCAGCCGACCC CGAAGACCCCGCGCCAACT GCGTGAGTTCCTGGGTAAA GCGGGTTTTTGCCGTCTGTT TATCCCGGGTTTCGCGGAAA TGGCGGCGCCGCTGTACCC GCTGACCAAACCGGGTACC CTGTTTAACTGGGGTCCGGA CCAGCAGAAAGCGTACCAA GAGATCAAACAGGCGCTGC TGACCGCGCCGGCGCTGGG TCTGCCGGACCTGACCAAG CCGTTCGAGCTGTTTGTTGA TGAAAAGCAGGGTTATGCGA AAGGCGTTCTGACCCAGAAA CTGGGTCCGTGGCGCCGTC CGGTTGCGTACCTGAGCAA GAAACTGGATCCGGTTGCG GCGGGCTGGCCGCCGTGCC TGCGTATGGTTGCGGCGAT CGCGGTTCTGACCAAAGAC GCGGGCAAGCTGACCATGG GTCAACCGCTGGTGATTCTG GCGCCGCATGCGGTTGAAG CGCTGGTTAAGCAGCCGCC |

| | | |
|---|---|---|
| | GGACCGTTGGCTGAGCAAC<br>GCGCGTATGACCCACTATCA<br>AGCGCTGCTGCTGGATACC<br>GACCGTGTTCAGTTCGGTCC<br>GGTGGTTGCGCTGAACCCG<br>GCGACCCTGCTGCCGCTGC<br>CGGAGGAAGGTCTGCAGCA<br>TAACTGCCTGGACATTCTGG<br>CGGAGGCGCACGGTACCCG<br>TCCGGATCTGACCGACCAG<br>CCGCTGCCGGACGCGGATC<br>ACACCTGGTATACCGACGG<br>CAGCAGCCTGCTGCAAGAA<br>GGCCAGCGTAAGGCGGGTG<br>CGGCGGTTACCACCGAGAC<br>CGAAGTTATCTGGGCGAAA<br>GCGCTGCCGGCGGGTACCA<br>GCGCGCAGCGTGCGGAGCT<br>GATTGCGCTGACCCAAGCG<br>CTGAAAATGGCGGAGGGCA<br>AAAAGCTGAATGTTTATACC<br>GATAGCCGTTACGCGTTTGC<br>GACCGCGCACATCCATGGT<br>GAAATCTACCGTCGTCGTGG<br>TTGGCTGACCAGCGAAGGC<br>AAAGAAATCAAAAATAAGGA<br>CGAGATTCTGGCGCTGCTG<br>AAAGCGCTGTTCCTGCCGAA<br>ACGTCTGAGCATCATTCACT<br>GCCCGGGTCACCAGAAAGG<br>TCACAGCGCGGAGGCGCGT<br>GGTAATCGCATGGCGGATC<br>AAGCGGCGCGTAAAGCGGC<br>GATTACCGAAACCCCGGATA<br>CCAGCACCCTGCTGATTGAA<br>AATAGCAGCCCGTAA*TAACT*<br>*CGAGTAAGGATCTCC* | |
| An example (GFP_TAAin) of functional PEgRNA cassette | ttgacagctagctcagtcctaggtataata<br>ctagtCTTGTCACTACTCTGAC<br>CTAGTTTTAGAGCTAGAAAT<br>AGCAAGTTAAAATAAGGCTA<br>GTCCGTTATCAACTTGAAAA<br>AGTGGCACCGAGTCGGTGC<br>TTGAACACC***TTA***ATAGGTCA<br>GAGTAGTGAtttttt | Green: J23119 promoter<br>Red: 20-nt spacer targeting GFPplus coding sequence<br>Black: 76-nt gRNA scaffold<br>Blue: RTT of the 3 prime extension, the target TAA is shown in bold, italic<br>Orange: PBS of the 3 prime extension |

**Supplementary Table 4. Spacers and 3 prime extensions used in this study**

| PEgRNA | Space (5'-3') | 3 prime extension (5'-3') | RTT length (nt) | PBS length (nt) |
|---|---|---|---|---|
| PEgRNA_GFP_TAAin_T4 | CTTGTCACTACTCTGACCTA | TTAATAGGTCAGAGTAGTGA | 7 | 13 |
| PEgRNA_GFP_TAAin_T5 | CTTGTCACTACTCTGACCTA | CTTAATAGGTCAGAGTAGTGA | 8 | 13 |
| PEgRNA_GFP_TAAin_T8 | CTTGTCACTACTCTGACCTA | CACCTTAATAGGTCAGAGTAGTGA | 11 | 13 |
| PEgRNA_GFP_TAAin | CTTGTCACTACTCTGACCTA | TTGAACACCTTAATAGGTCAGAGTAGTGA | 16 | 13 |
| PEgRNA_GFP_TAAin_T20 | CTTGTCACTACTCTGACCTA | AAAAGCATTGAACACCTTAATAGGTCAGAGTAGTGA | 23 | 13 |
| PEgRNA_GFP_TAAin_PBS5 | CTTGTCACTACTCTGACCTA | TTGAACACCTTAATAGGTCAG | 16 | 5 |
| PEgRNA_GFP_TAAin_PBS8 | CTTGTCACTACTCTGACCTA | TTGAACACCTTAATAGGTCAGAGT | 16 | 8 |
| PEgRNA_GFP_TAAin_PBS17 | CTTGTCACTACTCTGACCTA | TTGAACACCTTAATAGGTCAGAGTAGTGACAAG | 16 | 17 |
| PEgRNA_GFP_T198A | CTTGTCACTACTCTGACCTA | TTGAACACCTTAGGTCAGAGTAGTGA | 13 | 13 |
| PEgRNA_GFP_T196C_T198C | CTTGTCACTACTCTGACCTA | TTGAACACCGTGGGTCAGAGTAGTGA | 13 | 13 |
| PEgRNA_GFP_198Tdel | CTTGTCACTACTCTGACCTA | ATTGAACACCTAGGTCAGAGTAGTGA | 13 | 13 |
| PEgRNA_GFP_combo | CTTGTCACTACTCTGACCTA | TTGAACACTTATTAGGTCAGAGTAGTGA | 15 | 13 |

| | | | | |
|---|---|---|---|---|
| PEgRNA_GFP_12in | CTTGTCACTACTCTGACCTA | TTGAACACCTGTACAGATTACATAGGTCAGAGTAGTGA | 25 | 13 |
| PEgRNA_GFP_18in | CTTGTCACTACTCTGACCTA | TTGAACACCCTATAGTGAGTCGTATTAATAGGTCAGAGTAGTGA | 31 | 13 |
| PEgRNA_GFP_33in | CTTGTCACTACTCTGACCTA | TTGAACACCCTCGCCCGCTCGGTTTCGTGGGCTGATTCTACAATAGGTCAGAGTAGTGA | 46 | 13 |
| PEgRNA_GFP_10del | CTTGTCACTACTCTGACCTA | GGGAAAAGCATTGGTCAGAGTAGTGA | 13 | 13 |
| PEgRNA_GFP_23del | CTTGTCACTACTCTGACCTA | TGATCCGGATAACGTCAGAGTAGTGA | 13 | 13 |
| PEgRNA_GFP_36del | CTTGTCACTACTCTGACCTA | ATGCCGTTTCATGGTCAGAGTAGTGA | 13 | 13 |
| PEgRNA_GFP_49del | CTTGTCACTACTCTGACCTA | TCTTGAAAAAGTCGTCAGAGTAGTGA | 13 | 13 |
| PEgRNA_GFP_97del | CTTGTCACTACTCTGACCTA | CTTTGAAAGATATGTCAGAGTAGTGA | 13 | 13 |
| PEgRNA_lacZ_TAGin | AATCCCGAATCTCTATCGTG | TCAACCACCTACGCACGATAGAGATTCGG | 16 | 13 |
| PEgRNA_lacZ_CGdel | TATGCAGCAACGAGACGTCA | ATGAGCGGCATTTTCTGACGTCTCGTTGCTGC | 18 | 14 |
| PEgRNA_lacZ_GTtoTAsub | GCGAGTTGCGTGACTACCTA | ACTGTTACCCGTTAGTAGTCACGCAACT | 14 | 14 |
| PEgRNA_galK_TAAin | GACAGCCACACCTTTGGGCA | GCAGTTTCCTAAATGCCCAAAGGTGTGGC | 16 | 13 |
| **nsgRNA** | **PE3 or PE3b**[1, 10] | **Space (5'-3')** | | |

| | | |
|---|---|---|
| nsgRNA_GFP_TAAin | PE3b | AAAGCATTGAACACCttaAT |
| nsgRNA_lacZ_TAGin | PE3b | CctaCGCACGATAGAGATTC |
| nsgRNA_lacZ_CGdel | PE3 | CTGGAGTGACGGCAGTTATC |
| nsgRNA_lacZ_GTtoTAsub | PE3 | CATTAAAGCGAGTGGCAACA |

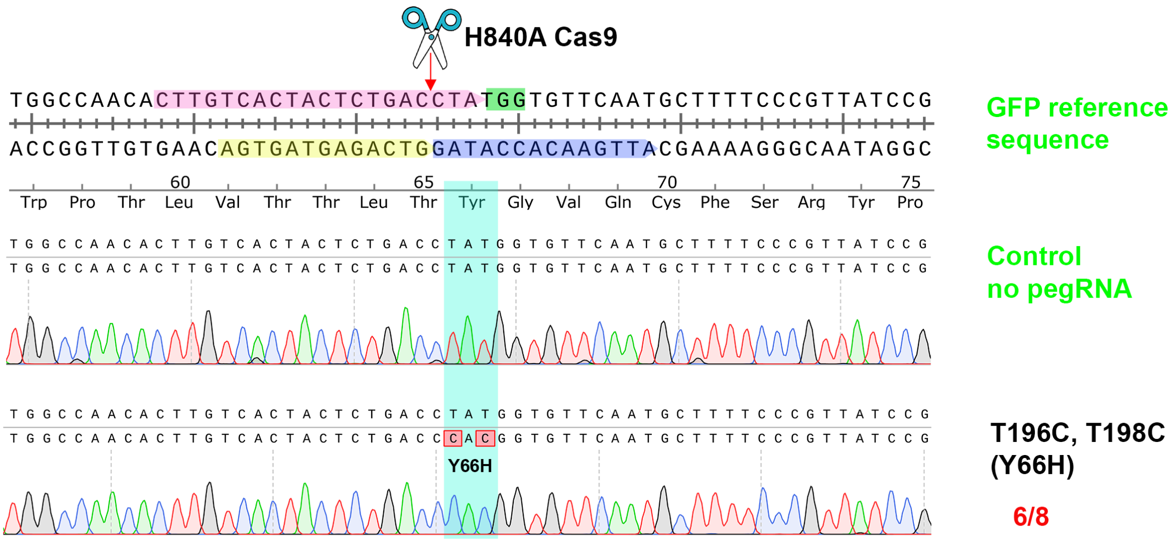

**Supplementary Fig. 1. A DNA editing of double substitutions by CRISPR-nRAGE.**

Eight randomly picked colonies of each designed DNA engineering were Sanger sequenced and traces were aligned to the targeted locus of GFP coding sequence. The correctly edited colony numbers and the total sequenced numbers were shown in red. The in-figure legend is the same as Fig. 2a.

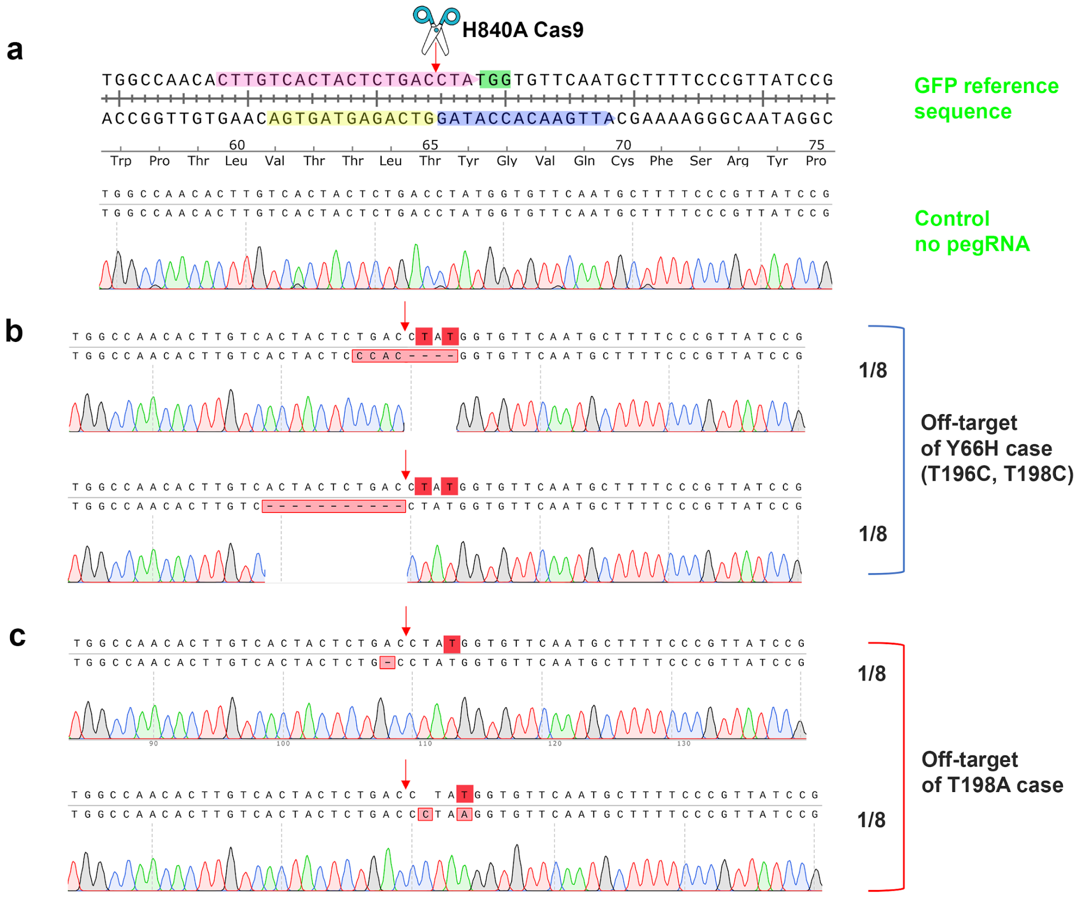

**Supplementary Fig. 2. Some observed target specific off-target effects of CRISPR-nRAGE for substitution.**

Eight randomly picked colonies of each designed DNA engineering were Sanger sequenced and traces were aligned to the targeted locus of GFP coding sequence. The incorrectly edited colony numbers and the total sequenced numbers were shown on the right of the trace. The nick site is indicated by a red arrow. The designed target nucleotide was highlighted by a red box. **a.** The in-figure legend is the same as Fig. 2a. A targeted GFP locus was displayed. **b.** Two clones with off-target were observed in the designed Y66H substitution case of in total 8 sequenced clones. **c.** Two clones with off-target were observed in the designed T to A substitution case of in total 8 sequenced clones.

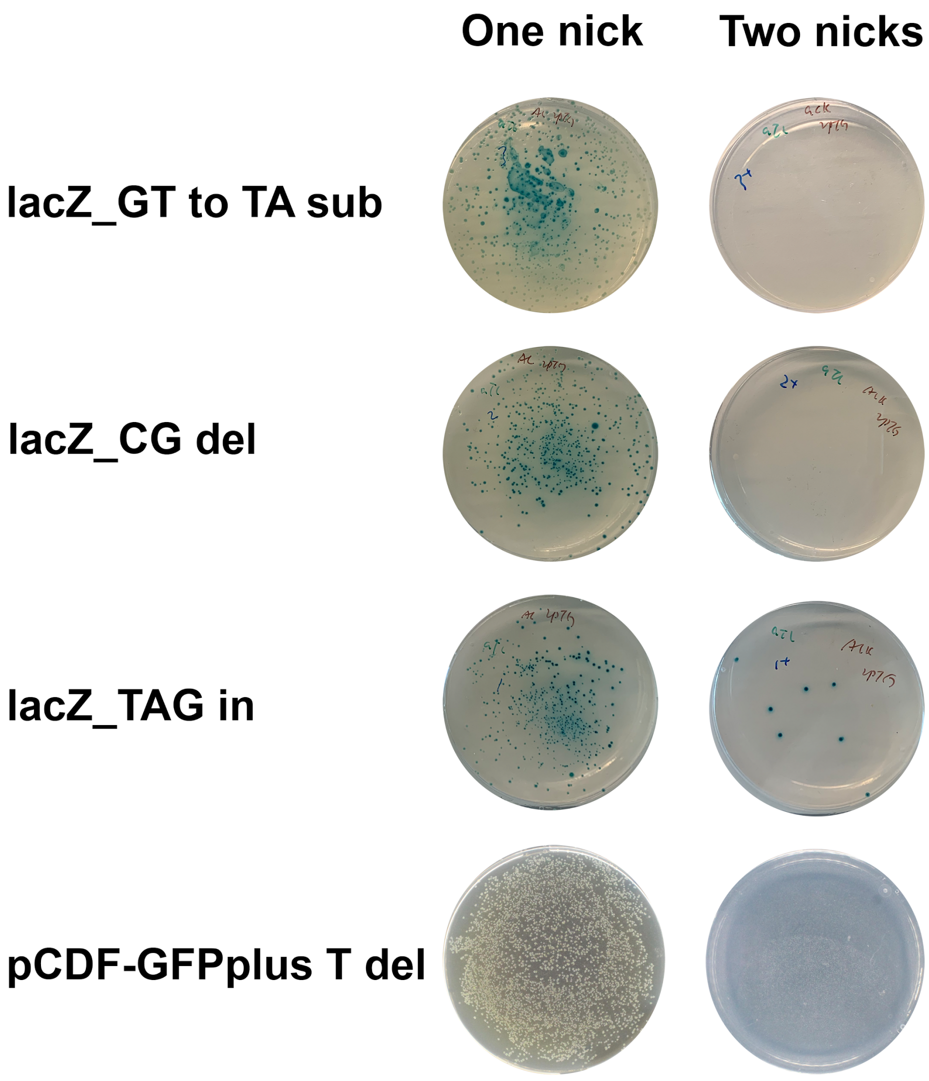

**Supplementary Fig. 3. A second nick compromises the application of CRISPR-nRAGE in *E. coli.***

Plates showed the colony formation of transformants. For the one nick panel, 50 ul of transformation culture was plated onto appropriate antibiotics supplemented LB plates, while for the two nicks panel, 400 ul of transformation culture was plated. Photos were taken by a Doc-It imaging station after 24h incubation at 37 °C.

**Supplementary reference:**

1. Anzalone, A.V. et al. Search-and-replace genome editing without double-strand breaks or donor DNA. *Nature* **576**, 149-157 (2019).
2. Lin, Q. et al. Prime genome editing in rice and wheat. *Nat Biotechnol* **38**, 582-585 (2020).
3. Mali, P. et al. CAS9 transcriptional activators for target specificity screening and paired nickases for cooperative genome engineering. *Nat Biotechnol* **31**, 833-838 (2013).
4. Ran, F.A. et al. Double nicking by RNA-guided CRISPR Cas9 for enhanced genome editing specificity. *Cell* **154**, 1380-1389 (2013).
5. Bowater, R. & Doherty, A.J. Making ends meet: repairing breaks in bacterial DNA by non-homologous end-joining. *PLoS Genet* **2**, e8 (2006).
6. Shuman, S. & Glickman, M.S. Bacterial DNA repair by non-homologous end joining. *Nat Rev Microbiol* **5**, 852-861 (2007).
7. Tong, Y., Charusanti, P., Zhang, L., Weber, T. & Lee, S.Y. CRISPR-Cas9 Based Engineering of Actinomycetal Genomes. *ACS Synth. Biol.* **4**, 1020-1029 (2015).
8. Qi, L.S. et al. Repurposing CRISPR as an RNA-guided platform for sequence-specific control of gene expression. *Cell* **152**, 1173-1183 (2013).
9. Rhodius, V.A. et al. Design of orthogonal genetic switches based on a crosstalk map of σs, anti-σs, and promoters. *Mol Syst Biol* **9**, 702 (2013).
10. Hsu, J.Y. et al. PrimeDesign software for rapid and simplified design of prime editing guide RNAs.
